# Subtype-Selective Neuronal Vulnerability and Compensatory Neuroglial Remodeling in Human Cortical Dementia

**DOI:** 10.64898/2026.10.03.756274

**Authors:** Bhaskar Roy, Prasun K. Roy

## Abstract

Whether the nodal vulnerability in dementia is solely represented by changes in cellular abundance or attrition of cellular transcriptomic landscape attribute to that transition in aging brain is yet to uncover with minutiae. To understand this complication, we have analyzed two snRNA-seq data specific to (I) dorsolateral prefrontal cortex (DLPFC): A comprehensive study of cellular profiling among cognitively normal and pathological ageing and dementia (AD) brain, (II) occipital cortex (OC) in the context of Nasu Hakola disease along with a tissue level bulk RNA-seq data corresponds to temporal cortex. We pinpoint cellular subpopulations their proportional changes within the nodal points and their transcriptomic profiling.

In DLPFC we observed alteration of mitochondrial and synaptic maintenance program in the pathological manifestation of disease. In OC we noticed depleted upper layer CUX2 neuron, PVALB and SST neurons on the contrary an enrichment of RORB and LAMP5 neurons with compromised metabolic states in the disease context. Glial populations are marked by representation of fibrous astrocyte, neuronal-supporting subpopulations associated with extracellular matrix remodeling programs. Proportional increment of intermediate myelinating oligodendrocyte and a decrease in mature myelinating subpopulation highlighted a compromised myelin repair attempt. In temporal cortex at tissue level we observed transcriptomic landscape adopted a neuroglial phenotypes in dementia brain. Cell-Chat analysis demonstrated a shift in cellular coordination towards glial population from neuron in NHD brain. A single cell-based drug reprofiling approach identified efatutazone and balaglitazone as broad-spectrum candidates for major cell types in occipital cortex, whereas major actionable biological targets belong to CDK subtypes and proteasome subunits. Overall, our findings imply cellular persistence do not necessarily substantiate molecular preservation, also the coexistence of cellular vulnerability alongside selective remodeling features, present a cellular framework on how to cope up with degenerative changes.

## Introduction

Ageing imparts multifaceted changes within the brain starting from macroscopic anatomical variables such as gray matter and white matter volume change, cerebral atrophy, ventricular hypertrophy to microscopic changes like Aβ neurofibrillary tangles accumulation, tau-hyperphosphorylation, lipofuscin aggregates (1,2,3,4,5). The severity of this pathological burden fluctuates person to person, as evidenced by a cognitively healthy individual with substantial amyloid load, which would otherwise be detrimental to an individual with the same magnitude of the pathological hallmark, vis-à-vis termed as cognitive resilience (6,7,8,9). A trade-off between pathological aging and geriatric disorders like dementia is not only determined by the threshold of neuropathological signatures but also shaped by cellular competence to preserve functional and structural alteration in a specific brain lobe(10,11). However, the molecular and cellular facets that drive the transition from pathological aging to dementia are not well defined. Susceptibility of neurons at the transcriptional scale varies widely in response to Alzheimer’s disease; therefore, analyzing the selective vulnerability of neurons will be an ideal way to investigate these unidentified features. The majority of the studies consist of a comparative analysis between healthy and advanced dementia cohorts, which eventually subsides into an intermediate state of pathological aging. Our study aims to bridge that gap by a cross-sectional assessment of healthy, pathological-aging, and Alzheimer’s cohort to decipher the fate of different cellular states and their dynamic changes in the above-mentioned conditions. In that direction, the dorsolateral frontal (DLPC) cortex is highly relevant to address our queries.

DLPFC corresponds to Brodmann areas 9 and 46 an important lobe that acts as a hub of brain’s executive control network and facilitates objective-oriented actions, cognitive flexibility, long term memory encoding, coordinates with the autonomic nervous system to perform allosteric cardiovascular control and working memory (12,13,14). A number of diseases are associated with DLPFCalong with Alzheimer’s such as major depressive disorder, schizophrenia, substance use disorder, posttraumatic stress disorder, and alcohol use disorder (16,17,18,19,20). The major components that construct the DLPFC intricate neural circuits are clustered in layer 3, pyramidal neurons in this layer mediate the working memory function. Different groups of GABAergic neurons support the pyramidal neuron’s function. Basket cells of PV neurons exert an inhibitory control on the perisomatic region of pyramidal neurons.

Several pieces of evidence have also demonstrated that the interplay between these two groups of neurons might influence the outcome of gamma oscillation. SST (somatostatin) neuron another group of interneurons involved in distractor resistance for this pyramidal neuron by means of distance mediated inhibition of pyramidal neuron dendrites. A chandelier cell, a PV GABA inhibitory neuron, confers inhibitory regulation over the pyramidal cell axon. Cooperation of each and every individual component of this circuit is vital to orchestrate working memory in DLPFC. Human prefrontal cortex is also marked by the presence of higher proportional level of L2-3CUX2 (ACVR1/CTHSD7A) L2-L3 intra telencephalic subtype. These intra-telencephalic subtypes span layer 2-6 (L2-L6); notable populations include L5 extra-telencephalic (FEZF2), L6 corticothalamic (SYT6), and L5 −6 near-projecting (HTR2C) (21,22,23,24,25).

The occipital lobe is responsible for visual processing and object recognition. Functional impairments of this lobe are associated with several anomalies, such as bilateral complete hemianopia, Anton syndrome, agnosia, and occipital lobe epilepsy. The major subpopulations of cells cluster in this lobe are glutamatergic L6 IT Car3, GABAergic SST(NPY), Inhibitory LAMP5 (LHX6) and non-neuronal cells like Astrocyte (AQP4), Oligodendrocyte progenitor cell (PDGFRA), Oligodendrocyte (MOG), Microglia(C1QB) (26). Compromised state of TREM2-DAP12 signaling in microglia leads to the onset of Nasu-Hakola disease, an early-state dementia (27,28). The autosomal recessive disorder is predominantly microglia-centric, the fate of other neuronal and glial cell populations under this condition is not well elucidated. In our study, we mainly focus on the involvement of these cellular populations and how their interplay contributes to the disease in the occipital cortex. Besides, we delve into dissecting how transcriptomic dynamics in the neuronal population is frontal cortex contrive neuro resilience. Essentially, we also attempt to investigate whether remodeling of the preserved molecular (transcriptomic) features corroborates with the tissue homeostasis or denotes a fragmented response. In the end, we undertook an approach to map a small molecule library with cell-specific disease signatures that could potentially identify cell-specific and broad-spectrum agents with the potential to reverse the disease signatures associated with NHD in the occipital cortex.

## Methodology

### 2.1. snRNA-seq Analysis

#### 2.1.a. Preprocessing and quality control

We performed our analysis on snRNA-seq (single-nuclear RNA-seq) data: GSE243292 on dorsolateral prefrontal cortex acquired from postmortem human brain samples. A total of 15 subjects (CN=2,PA=5,AD=8) were included in this study, which consisted of normal brain, pathological aging, and Alzheimer’s disease human brain. Another snRNA-seq data on Nasu-Hakola Disease (NHD) subjects: GSE190013 obtained from occipital cortex(n=14, NHD=11,Control=3) for a separate study. (Raw barcoded gene expression metric-s were incorporated in Seurat 4.5.1 and quality control is included in at the single nucleus level. The nucleus is excluded if they fail to meet the following criteria: cells with <200 expressed or >10000 expressed genes and a mitochondrial transcript content >5%. Seurat’s “*NormalizeData*” is used in order to log normalize the expression values (scaling factor:10000). Subsequently, we employed variance stabilizing transformation methods “*FindVariableFeature”*, nfeatures:2000 to identify top 2000 highly variable genes (HVG).

The batch effect generated due to the donor difference, inter-sample variability, differences in sequencing run that might interfere with the quality of analysis is mitigated, implementing Seurat’s anchor-based integration framework. This approach consists of independent sample-wise/subject wise specific variable features (n=2000) identification, “*FindIntegrationAnchors”* (for GSE190013) based canonical correlation analysis (dims= 1:20), followed by data integration (*IntegrateData*, dim:1:20) and scaling of integrated matrix/assay “*ScaleData”* prior to dimesonal reduction. After that, on that integrated data, we conducted Principal Component Analysis (“*RUNPCA*”). As per variance contribution and our inspection on “Elbowplot,” we retained the first 20 principal components. Graph-based clustering was mediated by *FindNeighbours* (dims =1.20) with a resolution parameter of 0.1 or 0.2 *(FindClusters*). Two-dimensional visualization of clusters was generated by Uniform Manifold Approximation and Projection (UMAP) (dims: 1:20) using *RunUMAP*. For GSE243292 we performed “*Harmony”* based batch correction.

Cluster Annotation is performed based on the observation of canonical markers and their proportion on the Dotplot, Violinplot, and FeaturePlot. Clusters with a trace of doublets and mixed lineage signatures are excluded from further analysis. Prior to Cell-subtyping highly expressed genes specific to particular cell types are obtained by means of “*FindAllMarkers”* function. After subtype annotation condition specific identities are assigned and differential expression is performed based on Wilcox-rank sum test (parameters: i.logfc.threshold = 0.25, ii.min.pct=0.10, ii. test.use = “wilcox”). The genes significantly differentially expressed were identified as per the following parameters: i. Adjusted P value (Bonferroni correction<0.05), ii. |log2 fold change| > 0.25.

#### 2.1.b. Cell-specific subpopulation analysis

To estimate condition-specific alteration within major brain cell types we attempt to characterize cellular subpopulation/subclusters. A particular cell subtype from the annotated major subtypes is extracted using “*subset”* function. Then each annotated major subtype is subjected to re-clustering and preprocessing with the same approach as it was implemented on major subtypes. After that, within the individual sample, the proportion of that subtype is determined by dividing the enriched subpopulation by the corresponding major subtypes. The extent of this subpopulation enrichment is compared across the condition and represented by a boxplot. Transcriptionally distinct signatures within a cellular subpopulation correspond to different conditions are delineated using Wilcoxon rank-sum test in Seurat. Finally, the condition-specific differentially expressed genes (DEGs) account for transcriptional heterogeneity in that subpopulation are depicted through heatmap. Statistical assessment is performed using a Bonferroni adjustment for DEG analysis and a Benjamini-Hochberg adjustment for the enrichment study (statistically significant level is adjusted to P < 0.05).

### 2.2 Bulk RNA seq analysis of temporal cortex

In order to get a comprehensive overview of the temporal cortex, we extracted HTSeq-derived gene-level counts from the GSE261775 dataset. The data comprises two different types of subjects, Alzheimer’s disease (AD, n=44) and healthy control (HC, n=50). The raw count is derived through read.delim(), where the Ensembl gene identifier is kept as the raw name. Sample metadata is contrived by arranging columns into the aforementioned two groups (AD, HC), and the comparison was mediated using the function ∼ condition. Median-of -ratio is used to facilitate library size normalization that estimated size factors and corrected sequencing depth, RNA composition across the samples. Samples are filtered, and low-count genes are excluded ≤10 to mitigate the noise. Differential gene expressions across the condition is performed with DESeq2 (hypothesis testing is done here by Wald statistics. Log2 fold change estimation is performed by apeglm. Multiple hypothesis testing correction is applied using Benjamini-Hochberg false discovery rate (FDR). Genes with adjusted P-value <0.05 are further processed for downstream analysis. Ensembl gene identifiers are converted to official gene symbols and Entrez IDs by trimming the version suffix. Normalization is performed with a variance stabilizing transformation (VST) followed by PCA on the normalized data. The output of the analysis is represented through a high-resolution Volcano plot (Enhanced Volcano) with adjusted P value and Log-fold change. Heatmap of the top-expressed genes is represented through a heatmap (intensity is adjusted through z score). Cluster Profiler is used for functional enrichment analysis of significant genes. After that using Entrez gene identifier we performed the GO pathway enrichment (Benjamini-Hochberg correction, FDR<0.05).

### 2.3 Cell Chat based cell-cell communication

To enumerate cell-cell communication across major cell types in he occipital cortex, we calculated the intercellular communication strength using the cell chat method. Using “Cell type” metadata the cell identities are designated. For our analysis, we have selected oligodendrocyte, Astrocyte, Ex_Neuron, In_neuron, microglia, and OPC as per the Seurat-based ‘subset’. The human ligand receptor interaction database (cellchat DB.human) is appointed to corroborate valid ligand-receptor interactions among the relevant gene set within each cell type. We used “create cell chat” function to construct cell chat object. After that, we used the subsetData()function to selectively pick signaling-associated genes. Using the “identify overexpressed Genes” we selected overexpressed genes within each cell type then to trace the cognate ligand receptor pair within sender and receiver population we employed *identifyOverExpressedInteractions()*. Communication probability is evaluated using cell chat’s mass action-based model compute communprob() based on average gene expression across cell type. To mitigate low confidence interaction, we used “filterCommunication(min.cells = 10)” interactions prominent in at least 10 cells per population. Followed by determination of pathway level communication probability via compute commun prob pathway and construction of global intercellular network by means of aggregateNet(). NetAnalysis_compute Centrality () function determines network centrality metrics important to decipher dominant signal senders and receivers across pathways. Netvisual enables us to visualize global communication circles, whereas interaction strength is defined with netvisual_heatmap().

### 2.4 Reprofiling of potential therapeutic molecules based on single nuclear signatures

We performed a single-cell-based reprofiling of potential therapeutic candidates with a disease/pathology reversal potential in occipital cortex. We worked on a disease reversal framework from the CMap/LINC database at the Broad Institute. Drug perturbation data comprises cell type transcriptional reversal score, determined based on changes in the expression profile of genes in different cell types in response to drug treatment, unique perturbation ID (cmap_name) and reversal score (rev_score). We used pivot_longer() to make the resulting database more structurally coherent. After that we have integrated our DEG data from CNS cell types (Excitatory neuron, inhibitory neuron, Microglia, OPC, oligodendrocyte and astrocyte) with L1000 Drug-gene data by matching similar genes. For the similar gene-cell-drug signature, we enumerate the expression pattern of the gene (termed as direction) and calculate the reversal score as: - Drug Z × Direction. The threshold for reversal scores are as such: **Strong reversal ≤ -0.15≥ weak reversal**. After that, based on the reversal score, drugs were mapped to individual cells. We employed slice_max (), with_ties = FALSE, to construct the top “N” candidates per cell and represent them through a heatmap. Next, we also wanted to pinpoint CNS-specific small molecules that reverses disease specific signatures in the above-mentioned cell types for that purpose we used CNS specific keywords (GABA, dopamine, glutamate, receptor, ion channel, opioid, agonist, antagonist, reuptake and so on) relevant to Mechanism of actions (MOA) and target annotation. It was executed using stringr::str_detect (), if at least one MOA/target matches the keywords that are marked as CNS-specific molecules. These small molecules have multiple targets; to figure out how many drugs are specific to a target, we used: n_drugs. In order to identify most prominent targets we traced whether the distribution of the transcripts (class of transcripts) restricted to a particular cell or encompasses multiple cell types by employing: n_cells.

## 3. Results

### 3.a. Assement of dorsolateral pre frontal cortex cellular selective resilience in the pathologic progression of Alzheimer’s

As per our snRNA-seq data of human frontal cortex analysis we observed heterogeneity in neuronal subpopulation characterized by enrichment of 5 different neuronal subpopulations Deep CT, L6_CTh, IT_SATB, RELN_L2IT and Inhibitory neurons. Cell proportion analysis demarcated a neurodegenerative switch in neuronal subpopulation. Deep corticothalamic (Deep_CT) and L6 corticothalamic(L6_CTh) subpopulation showed higher abundance in the disease context whereas proportions of RELN-positive neuron and Inhibitory interneuron are reduced in disease context. Starting with L6_CTh subpopulation, which is the most abundant compared to other subpopulations across conditions (cognitive normal, pathological ageing and Alzheimer’s) in our study. They exhibit relatively higher proportions of Pathological ageing(n=8171) followed by AD (n=6027) and cognitive normal(n=1175). (Fig.1 A-G & S1)

**Figure 1:**
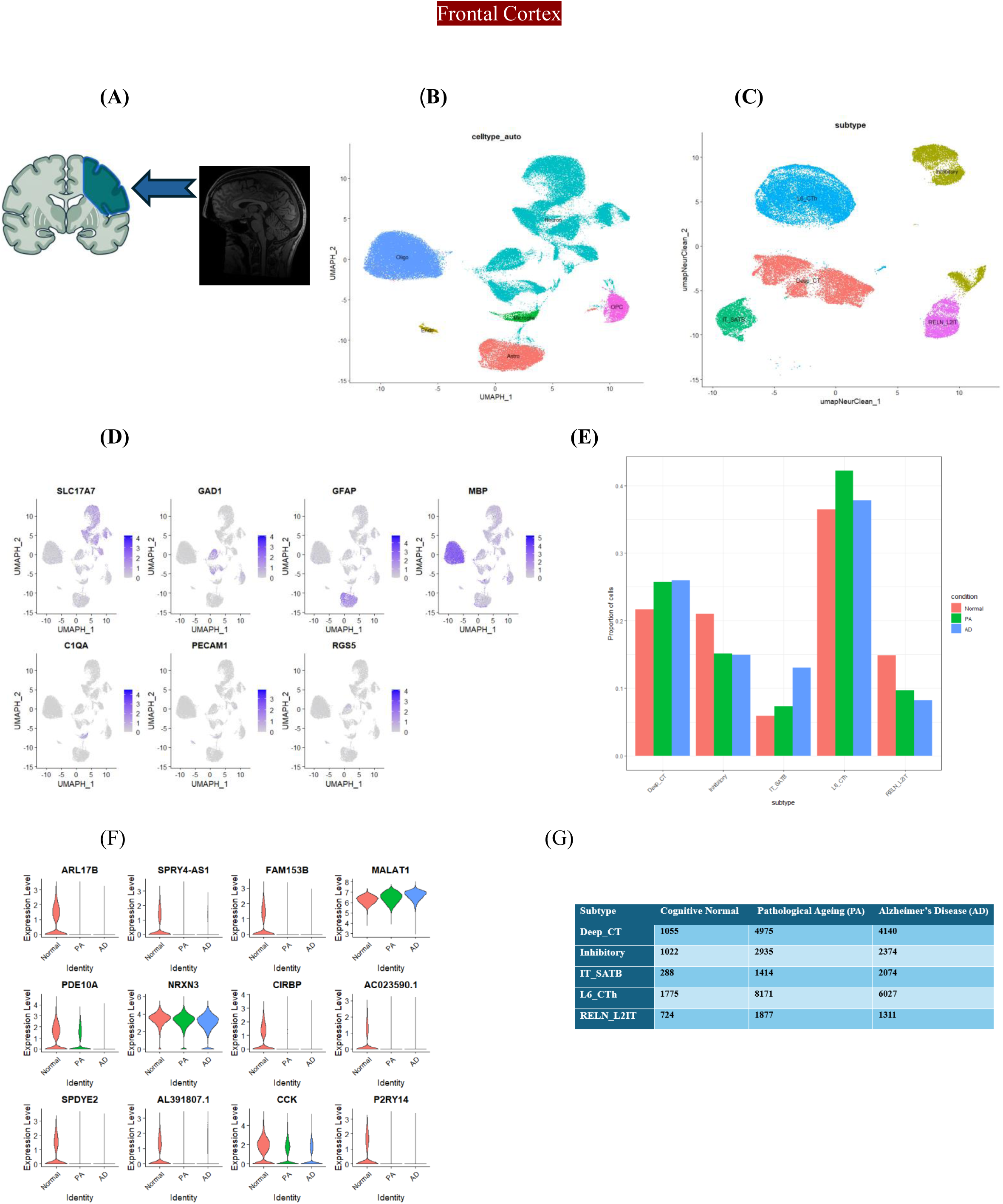
Cellular and Transcriptomic diversity in Dorsolateral Prefrontal cortex across Cognitive normal, pathological ageing and Alzheimer’s cohort. (A)We have analyzed the snRNA-seq data of dorsolateral prefrontal cortex (DLPFC) in CN, PA and AD;(B) Uniform Manifold Approximation and Projection (UMAP) of cellular diversity and (C) Neuronal subpopulation in DLPFC; (D) Distribution of cell type markers across major cell types (Neuron, Microglia, Oligodendrocyte, Astrocyte, OPC and Endothelial); (E) Violin plot represents proportional distribution of Neuronal subpopulations across different conditions. (F) Neuronal subtype markers expression across different conditions. (G) Abundance in nucleus number in Neuronal subpopulation in different cohorts.

The highly expressed transcripts (as per the heatmap) of L6_Cth subtypes fall into three major biological modules. The very first group comprises transcripts that play an important role in maintaining typical neuronal identity and signaling, such as P2RY14, ARL17B, SPRY4 -AS1, SNAP-AS1, FAM153B, SPDYE2, SPDYE5, and AC023590.1, which are enriched highly in the cognitively normal population and subsequently diminished with the expression within PA and AD (Fig. S3B) .

The next group comprises metabolic, mitochondrial module including MTCO2, MT-ND1, MTND4, MTATP6 involved in vital biochemical process like oxidative phosphorylation, ATP synthesis. GO term analysis showed a significant compromise with those vital functions, a clear indication of metabolic stress in the corticothalamic neuron. Since neuron is highly energy demanding and rely immensely upon mitochondrial energy currency (Fig. S3B,S2E).

The third major group of this analysis is the synaptic maintenance/neuronal projection module includes STMN1, RASGEF1B, LLRTM4, and NCAM2. We observed a sheer downregulation of these transcripts across all 3 cohorts. Since cortical neurons are deep projection neuron, a loss of transcripts actively involved in ATP production, metabolic homeostasis, neuronal extension, vesicle maturation account for a weakened corticothalamic regulation. On the other hand, an enrichment of transcripts associated with cell adhesion, synaptic organization, and calcium signaling suggests a compensatory approach to stabilize neuronal circuits, altogether an indication of bidirectional regulation(Fig. S3A).

#### 3.a.1. Deep_CT

In deep cortical neuron we observed enrichment of transcripts involved in synaptic plasticity such as NRX3, VGF, CCK, P2RY14, SPDYE2, CIRBP, ARL17B, FAM153B in cognitive normal and progressive attenuation in PA and AD. accompanied by reduced expression of mitochondrial transcripts such as NDUFA4, COX7A2, ATP51F and FTH1. As per the GO enrichment, we observed strong downregulation in Oxidative phosphorylation, ATP turnover, synaptic vesicle transport, and lysosomal cycling. Collectively these outputs depicts a reinstatement of neuronal circuit amid metabolic stress induced dysfunction(Fig.S3A&S2A).

#### 3.a.2. Inhibitory Neuron

Regarding the inhibitory neuronal population our analysis demarcated an enrichment of interneuron entity in the cognitive healthy cohort marked by the enrichment of SST, SCG5, PDE10A, ANKRD55, CIRBP, STMN1, BEX3, FTH1 followed by progressive loss and fragmented expression across PA and AD. Followed by the enrichment of mitochondria or energy metabolism transcripts like UQCRB,NDUFA4,COX7A2 are more heterogenous in disease. Notably, transcripts account for stress response and structural remodelling exhibit relatively consistent expression pattern across the conditions. Go pathway analysis also displays a similar pattern where downregulated genes are associated with oxidative phosphorylation, vesicle-mediated transport, proton motive force, and nucleotide biosynthesis. On the other hand, upregulated transcripts corroborated with the regulation of membrane potential, cell junction assembly, synaptic organization, and cell adhesion. The observed pattern signifies a potential compensatory reorganization to counteract the excitatory, inhibitory imbalance in dementia (Fig.S2D,S3D).

Additionally, a group level comparison between AD and CN cohort pinpointed upregulation of transcripts relevant to ECM remodeling (ADAMTS2) and RNA turnover (U2AF1,HNRLPLL,RALYL) a significant downregulation of stress response genes like CRH and CIRBP in AD,indicates a loss of CRH expressing neurons or its functional impairment (Fig.S1E).

Whereas a the comaparative anlysis between AD and PA display upregulation of transcripts such as LRRC63,CPNE2,DOCK10,CYP27A1,RFTN2; play crucial roles in calcium homeostasis,sterol homeostasis (S1F) in AD population.

### 3.b Systemic investigation of Occipital cortex cellular signature with a progressive age-wise alteration

We assess the cellular diversity in the occipital cortex of healthy and dementia brains (Nasu-Hakola-Disease) to trace the presence of any neurocognitive resilience signature(Fig.2A,2B). In the disease context, a prominent reduction in both excitatory and inhibitory neurons is observed, which indicates a global neuronal susceptibility during neurodegenerative changes. On the other hand, an enrichment of in the major glial population (astrocyte, oligodendrocyte, microglia and OPC) in dementia cohort is observed (Fig.2C). This coordinated cellular remodelling under neurodegenerative condition clearly reflects a systemic influence of neuro-glial population to offset the progressive impairment of neuronal function. It is estimated that a surge in overall glial population might buffer biological processes such as astrocyte, microglia-mediated metabolic homeostasis, immune surveillance, and synaptic reorganization, supported by enriched oligodendrocyte or OPC-mediated remyelination. This pattern underscores a neuro-resilience model where a compensatory response is possibly accrued by non-neuronal cells at tissue level to counterbalance deleterious and beneficial signatures(Fig.S4A,S4B).

**Figure 2:**
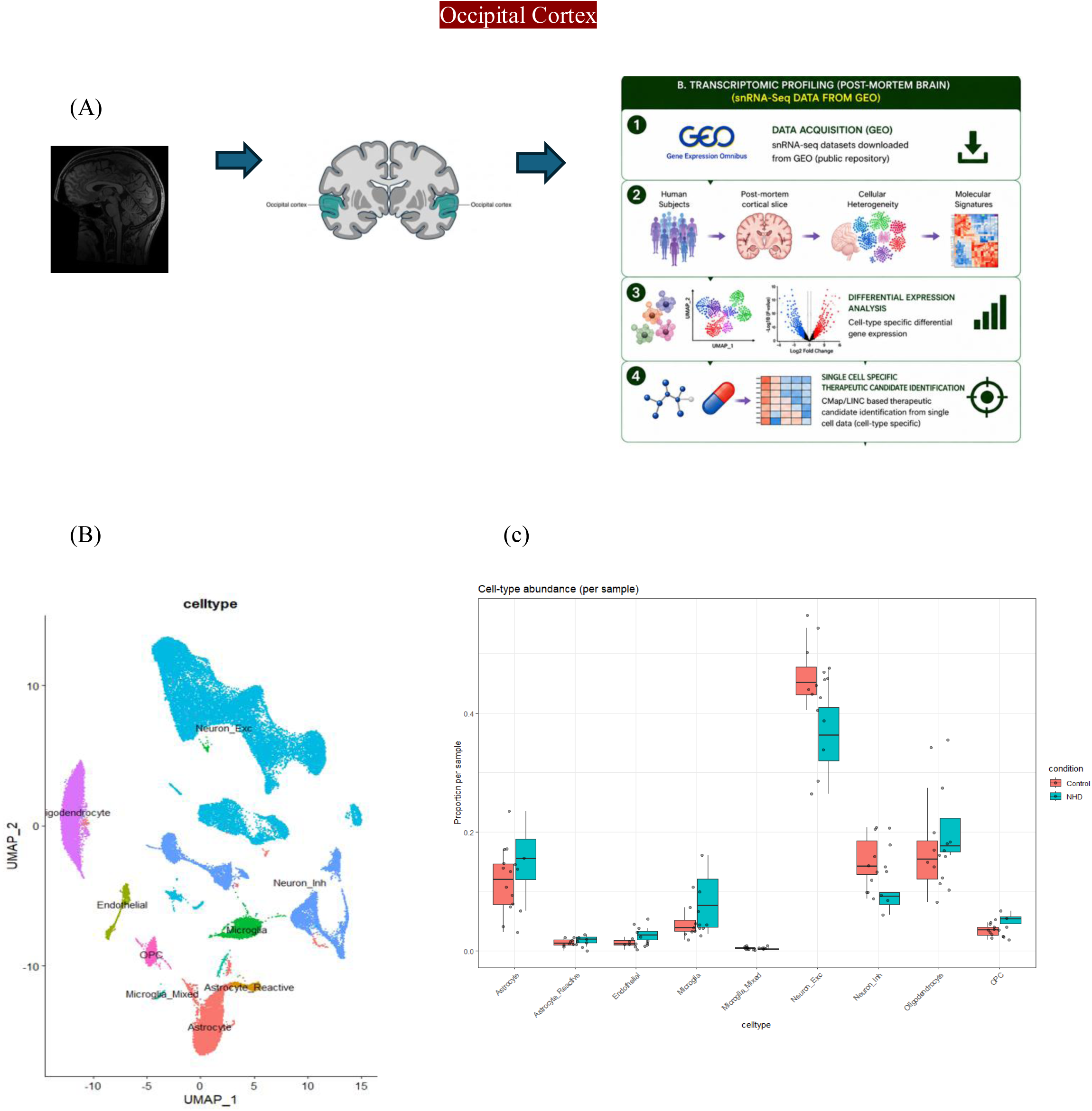
Cellular heterogeneity in occipital cortex (OC) in the realm of Nasu Hakola Disease (NHD). (A)Pipeline for occipital cortex snRNA-seq data analysis (Data acquisition, quality control, downstream analysis) (B)UMAP plot represents major cellular distribution in OC, (C) Proportion of major cell types of OC across conditions (Control vs NHD).

Further, our study aims to identify the condition-specific abundance of major cell-type subpopulations and underlying biological/functional traits to pinpoint potential gross protective measures undertaken by the brain.

#### 3.b.1. Excitatory neuronal populations exhibit a bimodal response

To comprehensively dissect the transcriptomic alteration in the occipital cortex under neurodegenerative changes we resolved its subpopulation. On a global level we observed a proportional increment of deep-layer RORB+ layer iv subpopulation of excitatory neurons in the disease context (NHD: Nasu Hakola disease). Notably, these neurons play a vital role in neocortex subtype identity in layer IV(EX_IT_L4_RORB). RORB-positive neurons turn out to be one of the early targets of tau pathology in the entorhinal cortex(Fig.3A,3B). However, we observed an increased proportion of excitatory intratelencephalic layer4 RORB neuron in dementia NHD cohort. In Alzheimer’s the RORB neuron is often characterized by loss of transcriptomic landscape associated with synaptic signaling. In the EX_IT_L4_RORB neuron a transcript level surge in healthy population is marked by enrichment of **EIF4A2**: translation initiation, **RPL3, RPL6, RPL14, RPL15, RPL32**: translation, ribosomal structural dynamics, **COX4I1**: oxidative phosphorylation, **PDP1**: pyruvate dehydrogenase, **PPIA**: protein homeostasis; **HSP90AA1, HSP90AB1**: stress response; in contrast, the diseased group is characterized by reduced expression of those above-mentioned transcripts (Fig.3C&3E). Together, these findings corroborate that transcriptomic signatures associated with translational landscape, mitochondrial plasticity and quality control systems are crucial for neuronal maintenance and their depletion leads to the onset of disease.Gene ontology analysis pinpointed upregulated biological processes related to synaptic organization, structural refinement, neuron projection development, dendrite development, and the synaptic vesicle cycle.

**Figure 3:**
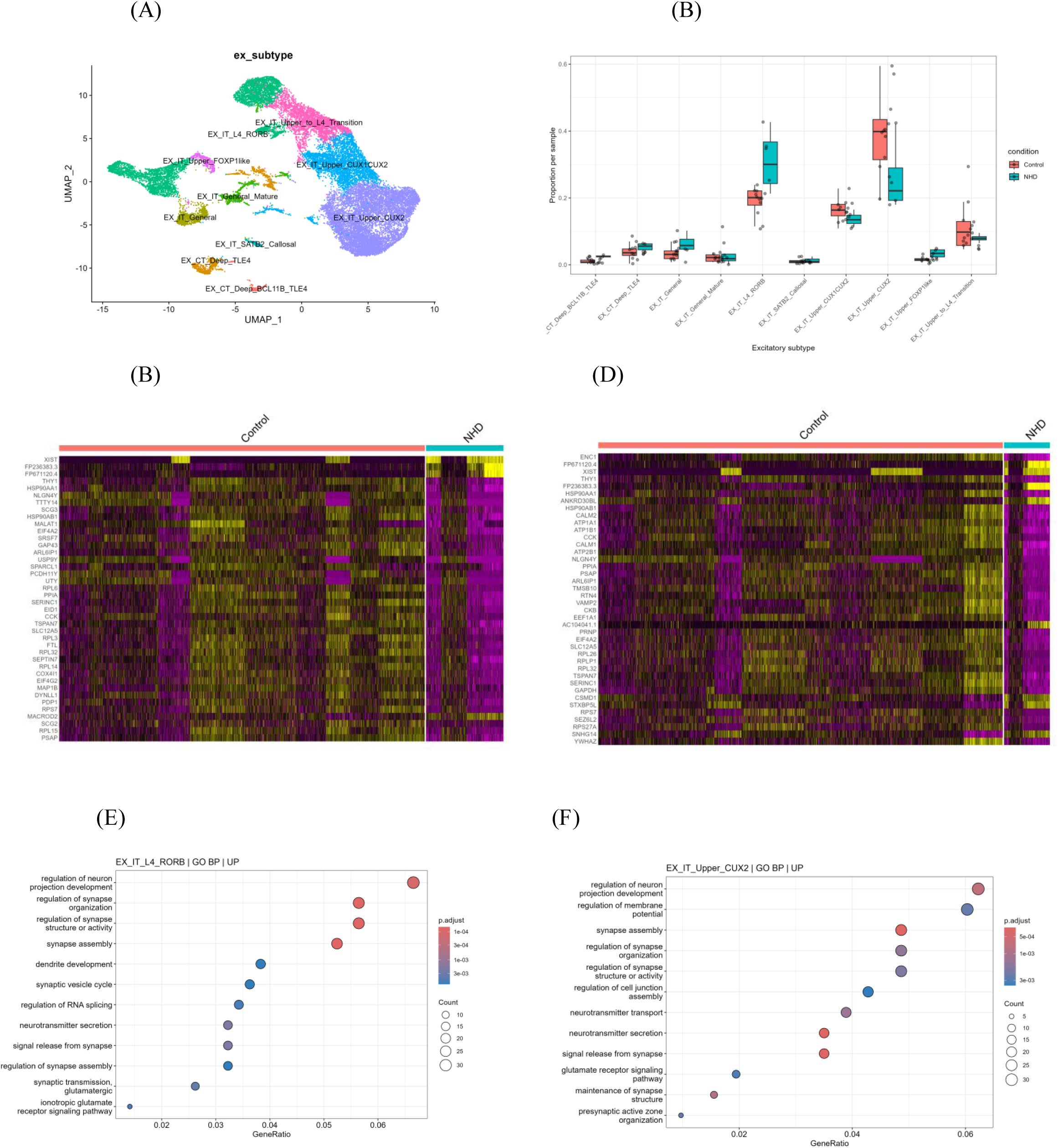
Transcriptomics landscape in OC Excitatory neuron. (A)UMAP representation of major excitatory neuron subpopulations distributed in OC. (B) Proportional changes of Excitatory neuron subtypes in different conditions (Control vs NHD). Heatmap denotes the enrichment of top genes in diseased (NHD) and Healthy (control cohort in (C) EX_L4_IT_RORB, (D) EX_IT_Upper_CUX2. (Bonferroni correction<0.05), |log2 fold change| > 0.25). (Upregulation is denoted by Yellow and downregulation by purple) GO enrichment of biological pathways based on upregulated transcripts in (E)EX_L4_IT_RORB, (F) EX_IT_Upper_CUX2 subpopulations (Benjamini-Hochberg correction, FDR<0.05).

Alongside, downregulated biological pathways are related to ribonucleoprotein complex biogenesis, ribosome biogenesis, translation, cellular respiration, and mitochondrial organization. The loss of translational regulation and mitochondrial maintenance in NHD EX_IT_L4_RORB neuron indicates an impaired energy supply as a of the result cell enduring a homeostatic strain.

We come across a significant number of transcripts that are enriched in disease cohort (NHD) associated with synaptic plasticity, axonal growth, neurotransmission, post-translational modification such as: **SPARCL1**(synaptic maintenance), **GAP43** (axonal growth), **TSPAN7**(dendrite spine formation), **SLC12A5/KCC2**(membrane potential), **MAP1B** and **SEPTIN7**(microtubule stability and cytoskeletal remodeling), **MACROD2**(post-translational modification), SCG3. The presence of synaptic remodeling signatures within RORB-positive excitatory neurons supports the fact that in spite of reduced cellular homeostasis cells are trying to undertake an adaptive strategy to cope up with a constrained cellular state (Fig.3D,3F).

Now our analysis demonstrated that Upper layer L2/3 neuron or precisely EX_IT_Upper_CUX2 neurons harbor a biphasic transcriptional reprogramming in diseased group where control subjects showed overall balanced expression. Major suppressed biological processes are translation, translation machinery, ribonucleoprotein complex biogenesis: RPL26,RPLP1,RPL32,RPS7,EEF1A1,EIF4A2;cellular respiration, mitochondrial gene expression and the associated transcripts: ATP1A1, ATP1B1,CKB; protein homeostasis: HSP90AA1,HSP90AB1. Preliminary we observed an augmentation in processes like neuronal projection(RTN4), synapse assembly, synaptic organization, neurotransmitter secretion: CSMD1, ANKRD30B, LYWHAZ, THY1, CKB, RTN4, TSPAN7 (Fig:S5A,SB)

#### 3.b.2. Persistance of inhibitory neuron subpopulation with compromised metabolic state

In the SST_inhibitory neuron population we noticed enrichment of signatures related to structural reorganization that is coupled with energy metabolism in the healthy cohort whereas, in the NHD cohort we noticed heterogenous and patchy expression of a significant numbers of transcripts associated with synaptic structure and mitochondrial functionality. Transcripts are associated with the process are CAMD2: regulates synaptic adhesion, SEZ6L2 and EXT1: dendrite outgrowth, neuronal maturation, NBEA: synaptic membrane adhesion and SCG3: secretory granule biogenesis, UBXN2B: nucleo-cytoplasmic shuttling, LRP8: synaptic-plasticity, RBFOX3, RPS6KA3: neuronal differentiation and neurogenesis, MT-ATP8, MT-ND1, MT-CO2: brain energy metabolism. Biological processes that are actively involved as per GO enrichment of IN_SST neuron are: neuron projection, regulation of membrane potential, cell-cell adhesion, synaptic organization, regulation of synaptic plasticity, post-synaptic potential, and axonogenesis. This might be an indication that the SST neuron entered into a metabolic compromised state highlighted by mitochondrial and synaptic functional perturbation that can actively reduce the gating fidelity of inhibitory neurons (Fig 4A,4B,4E & S5D).

**Figure 4:**
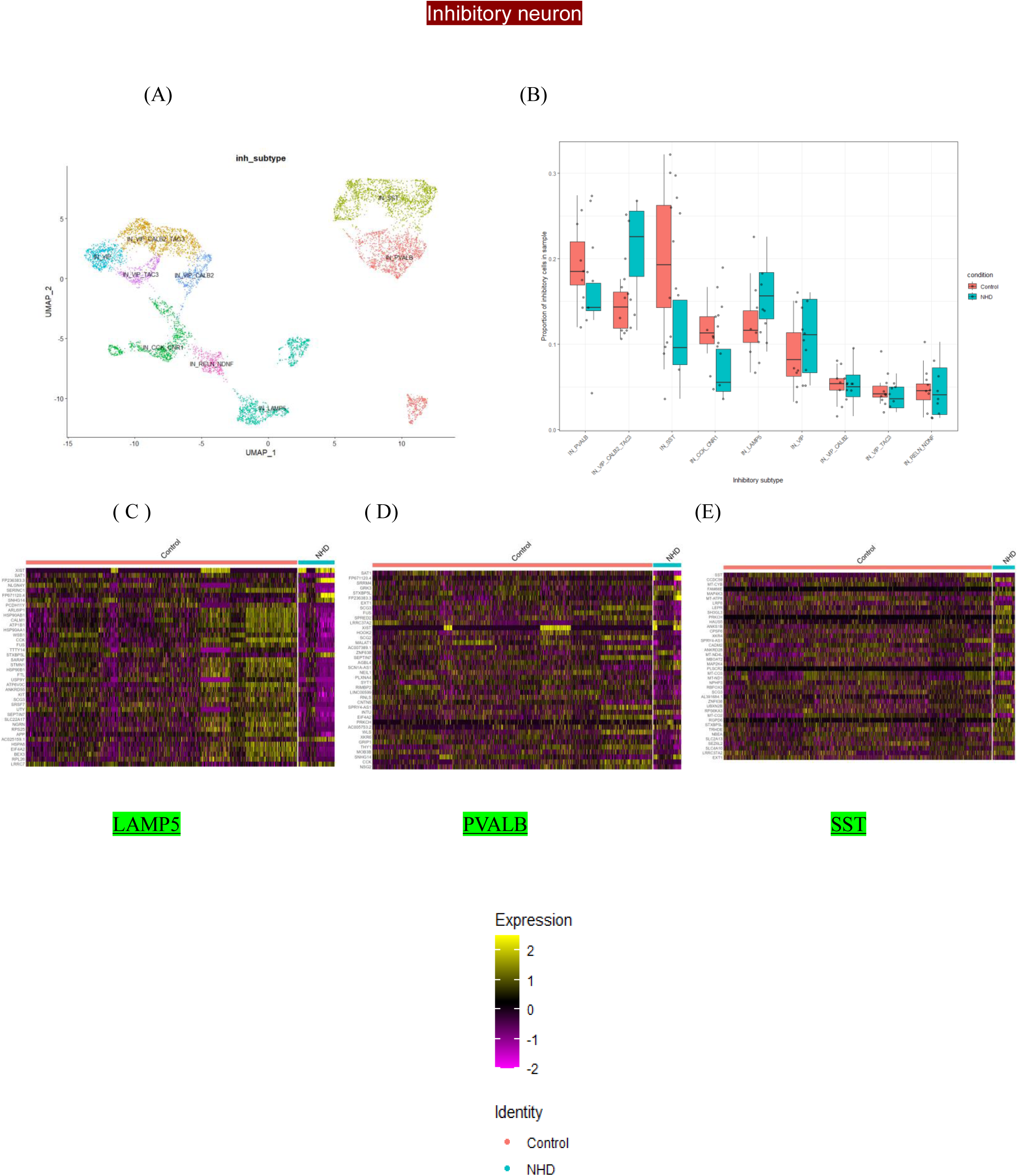
Transcriptomics diversity in OC Inhibitory neuron. (A)UMAP denotes inhibitory neuron distribution in OC;(B) Proportional alteration in major inhibitory neuron subpopulations in different healthy vs disease cohort. Heatmap shows the DEGs in (C) LAMP5 inhibitory neuron, (D) PVALB inhibitory neuron and (E) SST inhibitory neuron.

PVALB neuron, a GABAergic fast spiking neuron, proportionally decreased in diseased (NHD) cohort, with a median proportion reduction is ∼0.14 in NHD to ∼0.19 in control (∼25%). As per the adjusted p-value (<0.05) there are 260 DEGs in PVALB neuron including 192 upregulated in control and 68 in NHD. Transcripts with strong expressions in NHD are SCN1A-AS1, STXBP5L, GRK3, CNTN5, FP236383.3, XIST, LRRC37A2, INTU, FP671120.4. In control we observed enrichment of THY1, MT-ATP8, SRRM4, SCG2, SAT1, FXYD6, SCG3, FTL, HSP90AB1, SEPTIN7 (Fig 4B,4D).

Go enrichment analysis depicted the major biological processes involved are synapse structural organization, membrane potential regulation, potassium ion transport, calcium homeostasis, and dendrite development (Fig.S5C).

We noticed an enrichment of LAMP5 inhibitory neurons in the NHD population as compared to the control population. A total of statistically significant (adjusted p-value <0.05) 707 DEGs are observed, out of which 57 are upregulated and 650 are downregulated in the NHD cohort as compared to the control. Transcripts that are highly expressed in healthy cohort are NLGN4Y, PCDH11Y, TTTY14, UTY, CCK, RNLS, SCG2, RIMBP2, WLS, XKR6, GRIP1, MOB3B, CNTN5, associated with cell adhesion, synaptic development, neural plasticity, catecholamine metabolism, neuroendocrine secretion, synaptic organization, vesicular secretion, apoptosis, and neurogenesis (Fig.4C).

Interestingly, we have discerned upregulation of a number of noncoding RNAs in NHD such as XIST (ad), FP671120.4(AD), SNHG14, AC025159.1, FP236383.3. previously these are reported to involved in exacerbating dementia and other neurodegenerative diseases.

#### 3.b.3 Selective redistribution of astrocytic subtypes in early dementia

Proportional analysis of astrocyte subpopulations in two different conditions (control vs NHD) demonstrated refinement of a number of subpopulations in disease conditions such as Fibrous astrocyte (ASTRO_Fibrous_ECM): median proportion Control 20% vs NHD 23% (approx); Synaptic supporting neuron (ASTRO_Neuron Like_Doublet): median proportion Control 21% vs NHD 26% (approx) metabolically stressed astrocytes (ASTRO_Metabolic_Stress): control 29% vs NHD 31%. In turn reactive astrocyte (ASTRO_Reactive) proportionally decreased in diseased group (Control 24% vs NHD22%). Overall, we observed 9793 DEGs, out of them 3543 were upregulated in NHD and 6250 were upregulated in control (Fig.5A,5B,5C).

**Figure 5:**
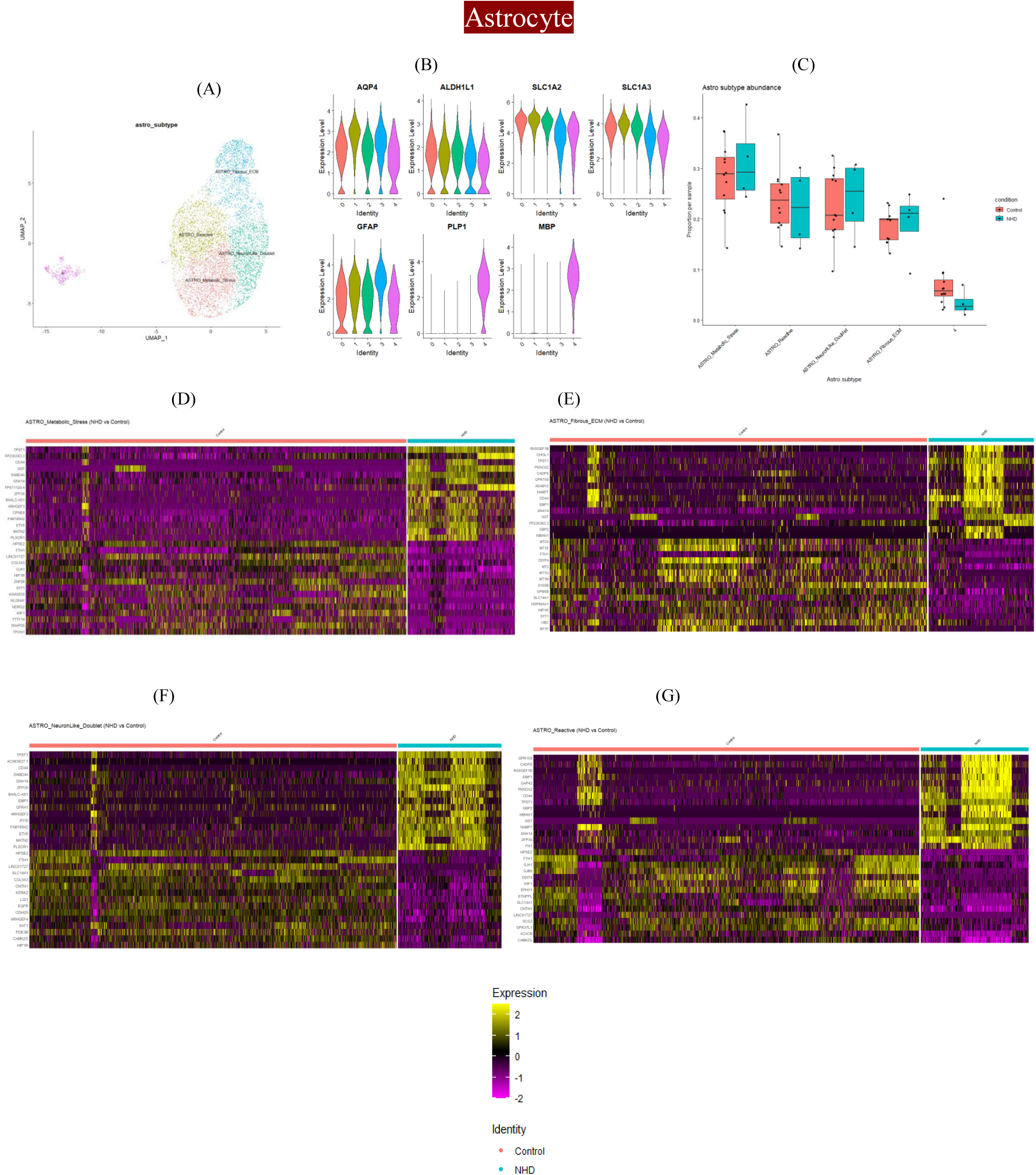
Astrocyte subtype and transcriptomic heterogeneity in remodeling OC microenvironment. (A)UMAP represents astrocyte subpopulation distribution in OC (B)Violin plot denotes the subtype specific markers expression (C) Barplot highlights the proportional distribution of astrocyte subtypes across different conditions. Heatmaps show the expression of top DEGs across different conditions (D) Astro_Metabolic stress, (E) Astro_Fibrous ECM, (F) Synaptic supporting neurons, (G) Astro_Reactive.

Within disease samples (NHD) in metabolically stressed astrocyte subtypes, we observed upregulations of transcripts associated with synaptic plasticity (TPST1), glial migration and differentiation (CD44), calcium homeostasis (CPNE8), membrane dynamics (PLSCR1), mRNA dynamics and translation (SAMD4A, ZFP36), and intracellular trafficking (ARHGEF3). In the healthy cohort we noticed an enrichment of transcripts associated with gap junction component (GJA1), Extracellular matrix remodelling (COL5A3), membrane trafficking (HIP1R), maturation of astrocyte (WIF1), iron homeostasis (FTH1), heparanase inhibitor (HPSE2), synaptic connectivity (NLGN4Y). GO pathway enrichment analysis revealed this subtype encompasses important biological processes such as development of neuronal projection, axonogenesis, cell-cell adhesion, small GTPase-mediated signal transduction, dendrite development, ameboidal cell migration indicating astrocyte’s attempt to remodel neuronal structure, respond to neuronal injury and maintain tissue homeostasis (Fig.5D,S6B).

In Fibrous Astrocytes (ASTRO_Fibrous_ECM) we noticed injury remodelling phenotype, as per the abundance plot, we could see there is a slight increment of this particular cell subtype-in NHD. The major transcripts that are enriched in NHD are involved in wound healing, cell substrate adhesion (RASGEF1B), cell matrix remodelling (EMP1, CHIL1), ameboidal type cell migration (CD44), inflammatory and cytokine responsive signaling (NAMPT), and small GTPase mediated signal transduction (GPR158, GNA14, GBP2)(Fig.5E&S6E).

In contrast within the control cohort we spotted a number of transcripts associated with neuroprotection and neural resilience such as metallothionines (MT1E, MT2A, MT1G, MT1M): stress response and metal buffering, S100B: confers protection against Aβ42, GPM6B: myelination, HSP90AA1: proteostasis, HIP1R: dendritic development. These finding indicates fibrous astrocyte is adapting a structural remodelling aimed at stabilizing damaged microenvironment and tissue integrity during neurodegenerative stress by promoting compensatory strategies such as astrocyte stress buffering, migration, cellular interaction and ECM remodelling. It is important to note that the persistent activation of this biological process might lead to a transition from protective to maladaptive glial scarring and a curtailed neural plasticity.

Synaptic supporting astrocyte (Astro_neuronlike) doublet proportionally increased in NHD as compared to control.Transcripts upregulated in normal brain showcased an elevated expression HPSE2,FTH1,SLC14A1,COL5A3,CNTN1,NTRK2, LGI1,EGFR,CDH20, ARHGEF4,SAT1. majorly involved in synaptic neuronal support. However, the diseased cohort represent an upregulation of a number of stress remodelling and inflammatory transcripts. That highlights even though this subpopulation is trying to preserve synaptic connectivity, neuronal communication, cellular adhesion but chronic neurodegenerative stress tends to foster an injury-associated transition (Fig.5F).

Reactive astrocyte subpopulation (ASTRO_Reactive), reported to act as a double-edged sword, proportionally depleted in the NHD cohort as compared to the control. In the diseased population we observed enrichment of transcripts such as PKNOX2, GAP43, CADPS, GPR158, RASGEF1B, EMP1, CD44, TPST1, GBP2, NIBAN1, XIST, NAMPTS associated with stress response, inflammatory signaling, axon growth and regeneration, vesicle docking, neural plasticity, chemotaxis, autophagy, neuronal differentiation, and cell substrate adhesion. As per the GO-down profile, we observed depletion of the aforementioned biological processes, which suggests a compromised state of synaptic homeostasis, organization, cellular communication, and maintenance of detoxification (Fig. 5G&S6C,S6D).

Whereas the control population exhibits enrichment of DDIT4, GPR37L1, ACACB, GJA1, GJB6, CNTN1, ETNPPL involved in cell substrate junction assembly, lipid homeostasis, cell substrate adhesion, neuroinflammatory response, ER stress this implies this subtype is trying to repair the neuronal milieu. Overall, our data illustrates that astrocyte subtypes are trying to maintain a stark balance between neural resilience and synaptic vulnerability.

#### 3.b.4. Oligodendrocyte population illustrated compensatory myelin remodelling but compromised bioenergetic states

Oligodendrocyte subpopulation analysis recounted a total of 1425 DEGs.We observed reduced mature myelinating (OLIGO_Mature_Myelinating) and increased myelinating intermediate (OLIGO_Myelinating_Intermediate) in the diseased (NHD) population as compared to control marked by a 14% median proportional decrease and 13% increase respectively (Fig.6A,6B,6C,6D).

**Figure 6:**
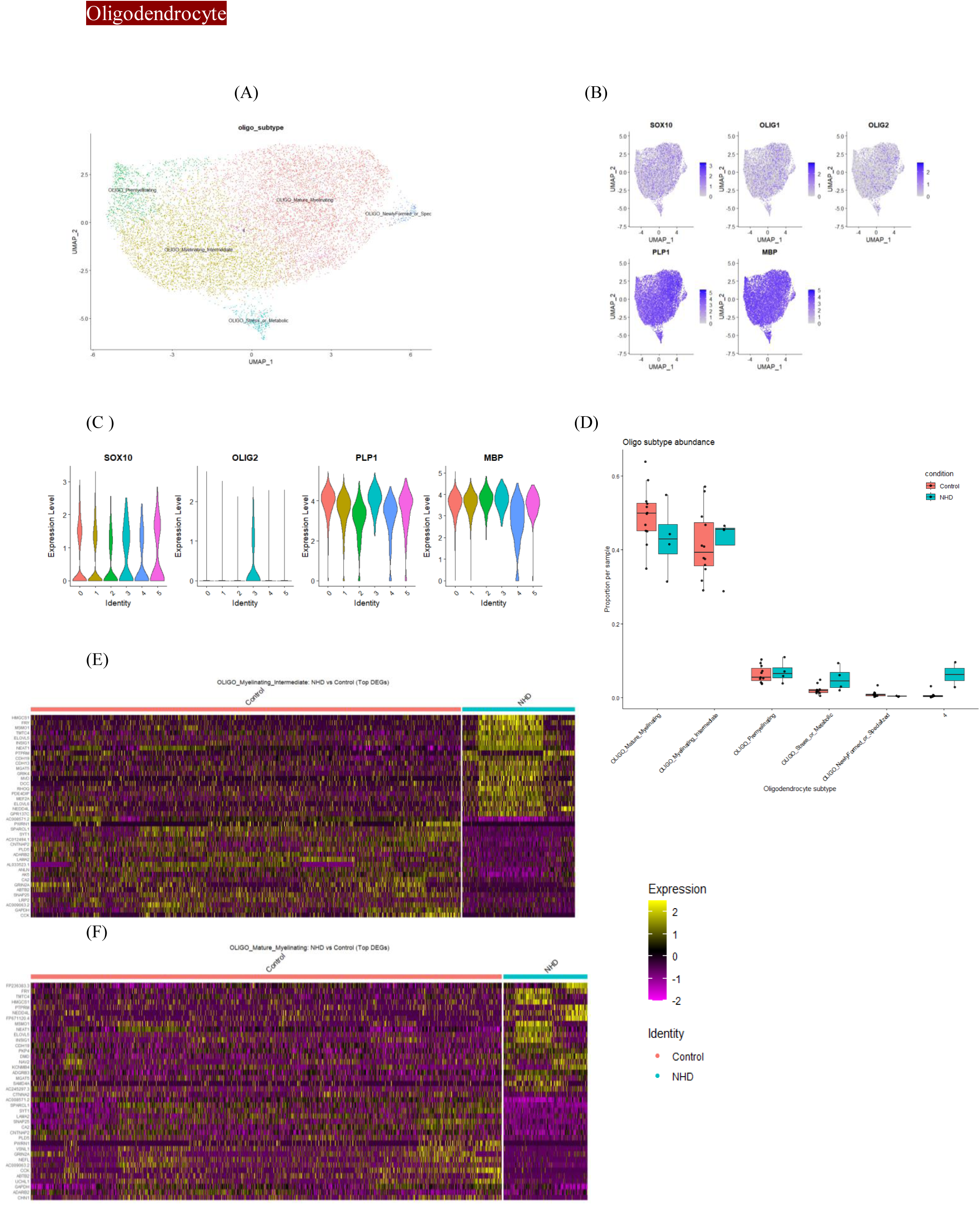
Oligodendrocyte transcriptional landscape in OC (A)UMAP of oligodendrocyte subtypes (B) UMAP of oligodendrocyte subtype as per classical markers distribution (C) Cluster-wise oligodendrocyte marker expression represented by violin plot. (D) Condition-wise oligodendrocyte subpopulation distribution across conditions. Heatmaps demonstrated top DEGs in (E) Oligo_Myelinating Intermediate and (F) Oligo_Mature Myelinating subtypes in Control vs NHD.

The transcriptomics of mature oligodendrocytes in NHD highlighted a re-myelination like program with a prominent enrichment of transcripts involved in cholesterol biosynthesis: HMGCS1, MSMO, post-translational modification and UPR: TMTC4, cell to cell adhesion:PTPRM, CDH19, long-chain polyunsaturated fatty acid synthesis: ELOVL5,OPC maturation:INSIG1,ADGRB3,cytoskeleton-dynamics:PKP4,NAV2,CTNNA2,ion buffering:KCNMB4,myelin-regeneration: MGAT5, oligodendrocyte differentiation:NEAT1 (Fig.6F).

In the healthy cohort, this subpopulation exhibits enrichment of more synaptic supporting signatures and (SPARCL1, CCK) vesicular trafficking (SYT1) alongside, oligodendrocyte maturation (LAMA2, GRIN2A) axonal projection, axogenesis and dendritic arborization (CTNAP2), neuroglial communication (GRIN2A), calcium homeostasis (VSNL1), protein folding (UCHL1), and myelin turnover (GRIN2A, CA2) (Fig.S9A,S9B).

Myelinating intermediate subpopulation shares a significant number of similar transcripts with the mature myelinating subpopulation additionally they express increasing level of MVD (cholesterol and steroid synthesis), DCC(axonal growth), RHOQ(neurite outgrowth), ANLN(cytoskeletal-dynamics), AK5 (neuronal energy metabolism),LRP2(Astrocyte-oligodendrocyte interaction), PDE4DIP(cAMP synthesis), MEF2A (lysosomal function, phagocytosis), GPRI137C (oligodendrocyte differentiation and myelination). Importantly, the set of DEGs in this subpopulation comprises a number of non-coding RNAs such as ACO12494.1, AC245297.3, AL0335231, AL033523.1 and a few noncoding RNAs shared by both subpopulation such as AC009063.2, AC008571.2. It will be interesting to study their proper physiological and functional implications especially in the realm of neuroprotective or neurodegenerative contexts (6E,S9C,S9D).

### 3.c. Cell-chat based cell-cell communication in occipital cortex

To elucidate the intercellular communication pattern in the occipital cortex we implemented a cell chat approach. We performed our analysis on six major cell types: oligodendrocyte, excitatory-neuron, inhibitory neuron, astrocyte, microglia, and oligodendrocyte progenitor cell. Cell chat, works on mass-action-based model and underscores molecular interaction propensity of a cell signaling pathway between two cells. To achieve that, we need to precisely recognize ligand or receptor enrichment in a particular cell type and simultaneously the overexpression of its cognate receptor or ligand in the other cell type. We estimated global communication strength by analyzing the average gene expression per cell group. Our study demonstrated the outgoing signaling pattern, which was majorly orchestrated by Microglia, followed by astrocytes and excitatory neurons. Neurexin (NRXN) turns out to be one of the major signaling molecule both in outgoing (astrocyte, microglia, excitatory neuron) and incoming signaling (astrocyte, microglia and oligodendrocyte) . Neurexin acts as a vital component of synaptic organization, synaptic communication and circuit maintenance whereas astrocytic NRXN plays an important role in long-term potentiation (LTP) and oligodendrocyte NRXN is pivotal for myelination (Fig. 7A–7C).

**Figure 7:**
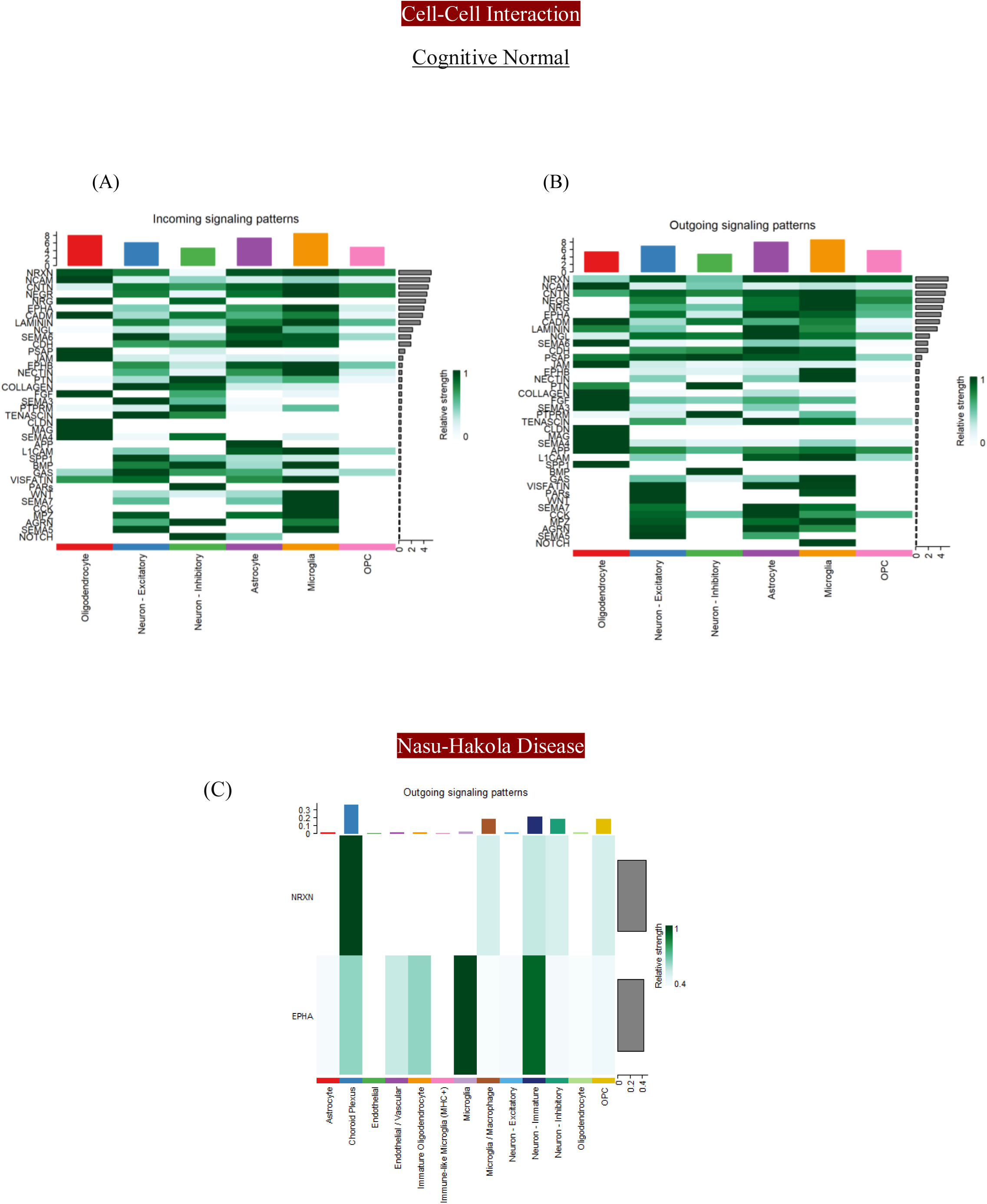
Heatmaps illustrated the interaction strengths among different major cell types in occipital cortex. (A) incoming signaling pathways denote receptors and (B) outgoing signaling pathways account for ligands. In healthy cohort. Whereas (C) NHD cohort is marked by only outgoing signals. Grey bar on the right stands for signaling strength of that pathway across cell types.

We also observed overlapping in a number of major signaling pathways among different cell types, such as Contactin (CNTN), which facilitates saltatory conduction and crafts the architecture of the node of Ranvier, shared by excitatory neurons, astrocytes, and neurons. Neuregulin (NRG): promotes neuronal survival by inhibiting apoptosis and inflammation, which is enriched in microglia, astrocytes (outgoing), and oligodendrocytes, inhibitory neurons (incoming).Ephrin type A receptor (EPHA): important in genesis and trajectory of nerve fibers, prominently shared by astrocyte microglia and excitatory neuron. Laminin: Blood-brain barrier integrity, neuronal outgrowth exerts its coordinated expression in astrocyte, oligodendrocyte,microglia (outgoing), and astrocyte,microglia,excitatory neuron (incoming).Cadherin(CDH): cellular survival, axonal projection, excitatory-inhibitory balance shared strong expression across excitatory and inhibitory neuron, astrocyte, microglia.

Our analysis also highlighted a number of fundamental developmental pathways exhibit a strong neuro-glial coordination like: Notch (inhibitory neuron -astrocyte), WNT(Microglia-Astrocyte-excitatory neuron), BPM (Neuron and microglia). Amyloid precursor protein (APP) that promotes the biosynthesis of amyloid beta is enriched in all the cell types.

Circle plot of intercellular networks also complements these findings. We observed a strong interaction from microglia to excitatory neuron, inhibitory neuron and astrocyte. In contrast, the net cellular interaction in the disease population discerned a collapse in that coherent network. Microglia lost most of its interaction with the glial population (oligodendrocyte, astrocyte), and excitatory neurons; the only major interaction is observed between microglia and inhibitory neurons (Fig. S7A,S7B).

### 3.d. Single-cell based potential therapeutic candidate for occipital cortex

Now, using a computational approach, we tried to identify small molecules with the potential to reverse cell-type-specific disease signatures in the occipital cortex. We map cell type specific DEGs of all seven cell types (Excitatory Neuron, Inhibitory Neuron, Astrocyte, Microglia, OPC, Oligodendrocytes and Endothelial) with perturbagen data (consists of FDA approved drugs) obtained from CMap/LINC database. The perturbagens that demonstrated strong disease reversal potential in given cell types are categorized as strong therapeutic candidates (P-value adjusted < 0.05, effect size cut off |avg_log2FC| ≥ 1). Drugs with a more negative reversal score signify how strongly that particular agent has the potential to reverse the disease signature of a given cell. The heatmap comprises the top 20 small molecules that can be potentially repurposed to treat dementia in the future.

Top candidate molecules exert their strong disease reversal features towards various cell types rather than being specific to one particular cell type. Most importantly, the top candidate molecules belong to several drug class such as triptolide (anti-inflammatory, immunosuppressive), OTS167(MELK inhibitor), CGP60474 (anti-infammatory), dinaciclib (antineoplastic agent/CDK inhibitor)(Fig.8B).

**Figure 8:**
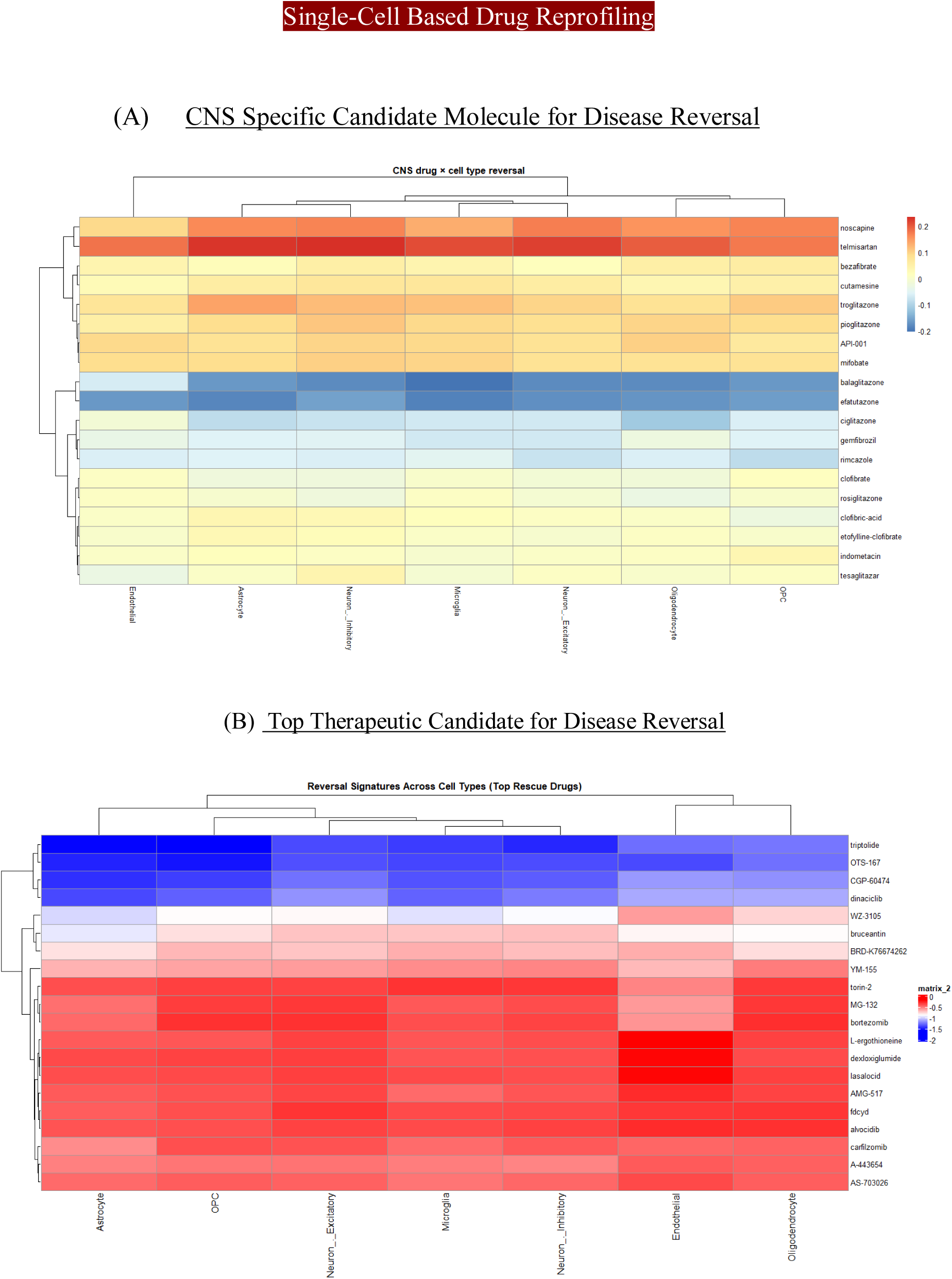
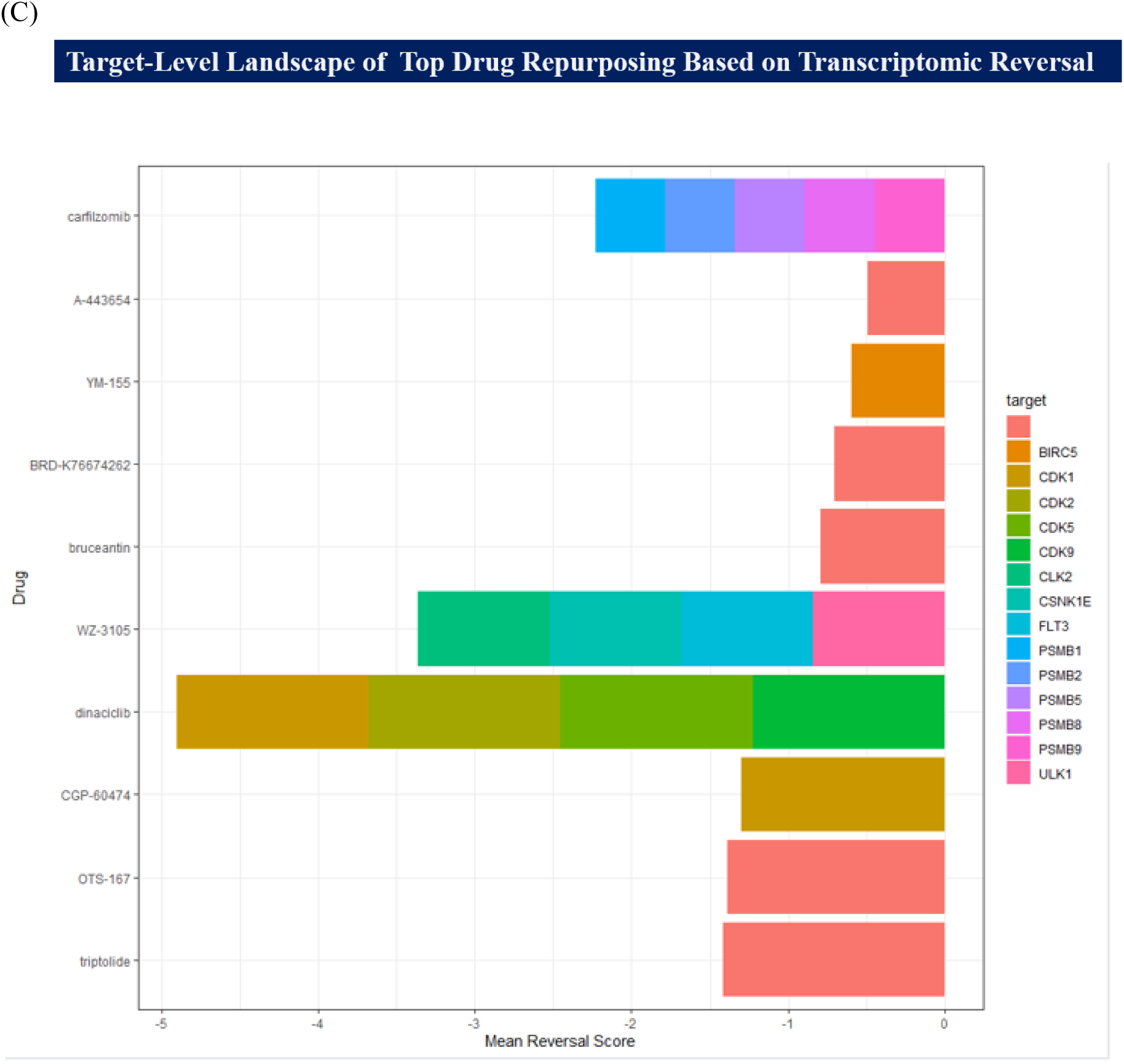
Cell specific identification of potential therapeutic candidates to reverse disease specific transcriptomic signatures per cell type in occipital cortex. (A)Central nervous system (CNS) specific potential small molecules. Efatutazone, Balaglitazone turns out to be broad spectrum potential to reverse the disease across major cell types. (Marked by blue in heatmap) (B) Top small molecules (Non-CNS specific) with a potential to reverse the signature across major cell types. (C) Top biological targets encompass CDK and proteosomal subunit classes.

Additionally, we strived to enumerate how the drugs of other CNS disorders could reverse disease signatures of dementia. The most significant drugs are efatutazone, balaglitazone, ciglitazone, gemfibrogil, Ciglitazone. The most prominent actionable drug targets belong to proteasome subunit and cell-cycle/CDK axis. This is an indicator that cells attempt to cope with the stress-coupled regulatory mechanism spanning the neuron and glial population (Fig.8B,8C).

### 3.e. Temporal cortex adopts a neuroglial remodeling at tisseue level

Whether our findings on neuroglial modulation at single-nuclear resolution reflect at the tissue level in a different lobe (other than frontal-occipital cortex), we estimate that dissecting bulk RNA-seq specific to the temporal cortex in the Alzheimer’s Disease (AD) context. We observed a distinct alteration of transcripts in AD cohort as compared to Control. Volcano plot shows a noticeable upregulation of DEGs associated with myelination:CD82, neuronal differentiation: NWD, Calcium influx: TRPV3, cognitive improvement, Potassium buffering: COL27A1, and myelination: ANLN (Fig.S8A). GO analysis highlights the major biological process that are enriched are axonogenesis, gliogenesis, ensheathment of neuron, myelination, oligodendrocyte and glial differentiation. Additionally, enrichment of astro-glial stress associated factors like KCNJ2, NEAT in AD cohort is also observed. Together, these findings in the temporal cortex intersect the central theme of neuroglial remodeling as observed in the occipital cortex. However, this tissue level bulk-RNA analysis doesn’t necessarily ensure a repair(myelin) mechanism, that require further molecular and physiological validation(Fig.S8B,S8C).

## 4. Discussion

Our study highlighted the regional heterogeneity of the cellular transcriptome along the dorsolateral prefrontal cortex and the occipital cortex, two ends of the dorsal stream of signal processing, in different grades of dementia compared to the healthy cohort. In the frontal cortex, we address the pathological vulnerability and associated resilience in Alzheimer’s disease and pathological aging. Whereas in the occipital cortex in Nasu Hakola disease, we observed neuronal population (excitatory and inhibitory neurons) succumbed to metabolic and translational paucity. But at the same time, multifaceted compensatory mechanisms were marked by the involvement of fibrous astrocyte, Oligodendrocyte mediated re-myelination and restructuring of other important neuro-glial extensions.

L6 corticothalamic neurons (L6CT) are specialized pyramidal cells in deep cerebral cortex, characterized by long range axonal projection and confer an excitation and inhibition through thalamocortical cells and thalamic reticular nucleus respectively. L6 neurons incorporate parvalbumin-positive interneurons to mitigate the metabolic stress and excitotoxicity generated by runaway glutamate release. They also the prevent the stroke or epilepsy associated hyper-synchronization (29,30,31,32). L6b excitatory and inhibitor neurons get activated through neuropeptide neurotensin and play a vital role in attention, wakefulness. Neurotensin is important in body temperature regulation as well as in neuroprotection; additionally, L6 neurons coordinate with microglia via MHC1 to maintain synaptic plasticity (33,34,35,36). Our findings demonstrated a bipartite status of the L6_CTh neuron. Loss of important synaptic maintenance/neuronal projection module STMN1, RASGEF1B, LLRTM4 and metabolic, mitochondrial module including MTCO2, MT-ND1, MTND4, MTATP6 that corroborate with synaptic structural loss and metabolic stress. Whereas, enrichment of transcripts like P2RY14, ARL17B, SPRY4-AS1, SNAP-AS1, FAM153B, SPDYE2 account for maintenance of neuroglial niche, neurite structure, synaptic adhesion, cytoskeletal rearrangement, and cellular differentiation. The combined findings depict an adaptive attempt to preserve the cortical circuit under metabolic and oxidative stress.

Our occipital cortex findings center around Nasu-Hakola disease (NHD), an autosomal recessive condition characterized by early onset of dementia (alongside bone abnormalities) and emerges due to mutations in TREM2, TYROBP, characterized by loss of myelin sheaths, severe fibrillary gliosis, atrophy in frontal and temporal lobes, axonal spheroids and basal ganglia calcification (37). In our analysis, we observed relative enrichment of L4_RORB neurons and depletion of upper-layer CUX2 neurons in NHD subjects. RORB neurons are marked by reduced expression of transcripts related to ribosomal structural dynamics, translation, and stress response; the impairments in these vital processes might lead to interference with synaptic protein homeostasis, axonal maintenance, and vesicular trafficking. The outcome indicates even though the RORB population is undergoing molecular dysfunction they have maintained relative cellular persistence. The RORB neuron in the diseased population also demonstrated an augmented expression of transcripts associated with synaptic plasticity, axonal growth, neurotransmission, membrane potential, and cytoskeletal dynamics. Cumulatively, these changes underlines RORB undergoes a metabolic and structural reorganization to minimize the drastic effect of reduced proteostatic stress. Whether these reorganizations are a cellular strategy to cope with the degenerative or injury-associated stress requires further spatial validation.

CUX2 neurons are excitatory projection neurons in the cerebral cortex upper layer II-III, important for cortico-cortical connections (38,39). CUX2 acts in base excision repair, and non-homologous end joining, as well as facilitates dendrite branching(40). The reduced relative abundance of CUX2 neurons in NHD subjects indicates a selective superficial disengagement of cortico-cortical circuitry, rather than uniform loss across a particular compartment. CUX2 neurons exhibit an enrichment in transcripts associated with synaptic and neuronal structural organization (TSPAN7, CSMD1, THY1, RTN4, TSPAN7). Concurrently, this neuronal population is also compromised with the vital biological processes like ribonucleoprotein complex biogenesis, cellular respiration, and protein homeostasis. The collective transcriptomic alteration in CUX2 neuron therefore accentuates the fact that even though the neuronal population is bioenergetically deficient and impaired proteostasis, they adopt a mode of structural reorganization to uphold the aforementioned changes. The result reconcile with the proposition that L2/3 neurons are selectively vulnerable in human Alzheimer’s disease. Previous findings from integrated single-cell study and spatial transcriptomics have pinpointed vulnerable L2/3 intratelencephalic neurons in supra-granular cortex and their association with excitatory neuron depletion (41). Our results therefore illustrate that the upper-layer CUX2 neuron might serve as a convergent point of vulnerability across different types of dementia.

In the healthy population, we observed proportional enrichment of parvalbumin interneuron (PVALB), Somatostatin-expressing neuron (SST). In contrast, there is a depletion in Lysosome-associated membrane protein5 (LAMP5) neuron, which implies NHD pathological manifestation is subtype specific. PVALB neuron is a fast-spiking GABAergic neuron and stabilizes Excitatory-inhibitory balance, regulate gamma waves and buffer calcium homeostasis. Lowered representation of this subpopulation in OC and impairments of major biological processes (synaptic structural organization, ionic flux, dendrite development) may interfere with the spatio-temporal efficiency of cortical inhibition (42,43,44). AD human studies have highlighted the enrichment of PV-neurons in the piriform cortex and a decline in the dentate gyrus (45,46). Whereas studies in the AD mouse model in the hippocampus demonstrated different levels of susceptibility of that particular subpopulation (47). Therefore, the exact mechanistic aspect of PVALB neuron in occipital cortex will be an interesting avenue to explore in future. SST neurons in layer 4 mediate the activation of pyramidal neurons and inhibit PVALB neurons (48,49). Previous research has demonstrated that SST deficiency is correlated with an increment in Aβ load. SST augments neprilysin levels, which are involved in Aβ degradation (50,51). A depletion in SST inhibitory neuron populations in NHD subjects implies impairment in dendritic responsiveness and excitatory post-synaptic potential. LAMP5 neuron proportionally increased in NHD populations compared to the control cohort. LAMP5 inhibitory neurons majorly reside in the superficial layer of the cerebral cortex (layer I, layer II, layer III) and play an important role in actively maintaining short-term synaptic plasticity, endosomal trafficking, and auditory gating. LAM5 is a presynaptic protein and loss of this protein compromises the theta-gamma coupling and impairs hypersynchronous neuronal activity. The loss of this inhibitory neuron population is evident in FTD and Alzheimer’s(52). Major pathology of the NHD centers around glial cells and the anomaly onset due to DAP12 - TREM2 signaling loss, the presence of the above-mentioned inhibitory neuron populations in the occipital cortex possibly aids in converging the feedforward signals with the sensory inputs that traverse through the superficial layers. Additionally, they might play a vital role in buffering neuronal hyperexcitability and synchronization of dendritic excitation. Overall, we noticed in disease condition, PVALB neurons exhibit the most prominent vulnerability, SST is marked by synaptic structural disruption and metabolic exhaustion, and LAMP5 neurons are relatively preserved, accounting for the cushioning of the occipital cortex circuitry. In the control cohort, conservation of the molecular program indicates the maintenance of a healthy brain aging hallmark. However, results in disease population represent an attenuated molecular states that corroborates with the neuronal vulnerability and functional inefficiency although more neurophysiological and molecular level validation is required in the support.

The glial population in the occipital cortex attempted to redistribute in order to withstand the neurodegenerative insults and myelin injury. One of the notable changes we observed here is the relative increment of metabolically stressed and fibrous/ECM-associated astrocytes, whereas the reactive population is depleted in the disease condition. Fibrous astrocytes harbor an injury remodeling phenotype related to cell matrix reorganization, cellular ameboidal type migration, and cellular adhesion. On the other hand, reactive astrocyte populations retain the signatures associated with stress response, arborization, vesicle docking, which accentuate even though cellular refinements are trying to mediate a structural and functional homeostasis to confer a coordinated neuroglial microenvironment, prolonged deposition of reactive subpopulation might elevate tissue scarring and undermine the synaptic plasticity.

Oligodendrocyte subpopulations are marked by an increase in the myelinating intermediate population and a reduction in mature myelinating groups. Mature oligodendrocyte in the disease cohort comprises signatures associated with cholesterol biosynthesis and fatty acid metabolism, possibly to reverse myelin loss. In the healthy population, they underpin oligodendrocyte maturation, myelin turnover, and synaptic support features. Myelin intermediate populations also illustrate neurorestorative phenotypes. Henceforth, these findings depict a possible involvement of oligodendrocytes in myelin restoration; concurrently, a reduction in mature oligodendrocytes may compromise the full remyelination potency.

Cell chat analysis underscores excitatory neuron as a major hub of cellular communication in healthy cohort. In the disease cohort we observed a fragmented cellular coordination pattern where microglia act as a hub of cellular communication. Microglial TGFβ1 axis regulates the facet of oligodendrocyte myelination, whether impaired TREM2 -DAP12 signaling (major root cause of NHD) might influence the incomplete oligodendrocyte mediated myelin repair process will be an interesting avenue to explore (53,27). In cell type specific perturbational candidate small molecule identification for CNS specific match we have pinpointed balaglitazone and efatutazone. Both of the molecules are reported to be PPARγ agonist, the first one plays an important role in lowering glucose level and the later in malignancies in colorectal and thyroid malignancies. PPARγ agonists selectively regulate the angiogenesis and inflammation through COX2, PGE2 pathways (57). Targeting PPARγ with agonist in astrocytes and microglia mitigate the amyloid loads and inflammation in mice model (58). Unrestricted screen help us to identify triptolide, OTS-167, CGP-60474, dinaciclib as top candidate with potential to reverse disease signatures across all cell types. Dinaciclib is earlier reported to inhibit the activity of CDK1, CDK2, CDK5 and CDK9 that is reconcile with our findings of actionable biological targets(60). However, further validation of these candidate small molecules in the specific cell line or animal model will give more transparent pictures on its actual neuroprotective/neurorestorative efficiency (61).

Our study aim to establish a mechanistic insight towards selective vulnerability in dementia across frontal, occipital and temporal cortex. We haven’t performed integrated analysis of the transcriptomic data form this three different nodes for a comparative evaluation, rather it is an empirical study to observe cellular perseverance amidst age-wise dementia associated neurodegenerative changes and transcriptomic rewiring. Unequal distribution of subjects from male and female population limited our study from a separate gender stratified analysis. However, our Cell-chat analysis and pinpointing major cell-type specific perturbational candidates provide a systematic avenue to focus on and prioritize specific cellular pathways, mechanisms and candidate molecule for further validation.

## Supplementary Figure

## Supplementary Figure

**S1.**
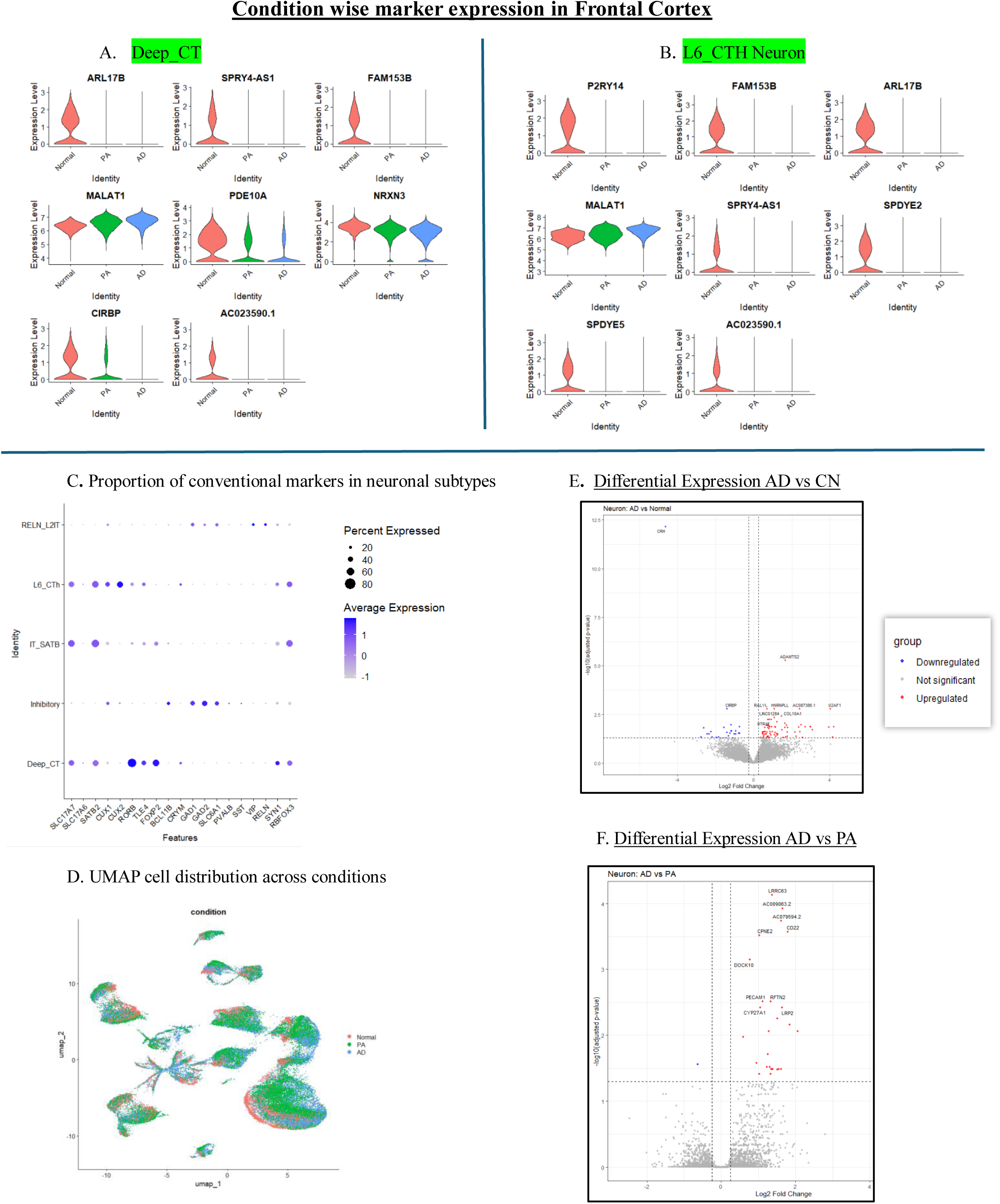
**A**. Violin plot represents the Cell type specific marker expressions in Cognitive normal, Pathological ageing and Alzheimer’s cohort in Deep corticothalamic neuron (Deep_CT) and (B) L6 corticothalamic neurons (L6_CT) (C) Dot plot represents abundance of markers in neuronal subpopulations in DLPFC. (D) UMAP accounts for condition-wise cellular (subtype) distribution. Volcano plot represents DEGs in (E) Alzheimer’s (AD) vs Cognitive Normal (CN), (F) Alzheimer’s (AD) VS Pathological ageing (PA).

**S2.**
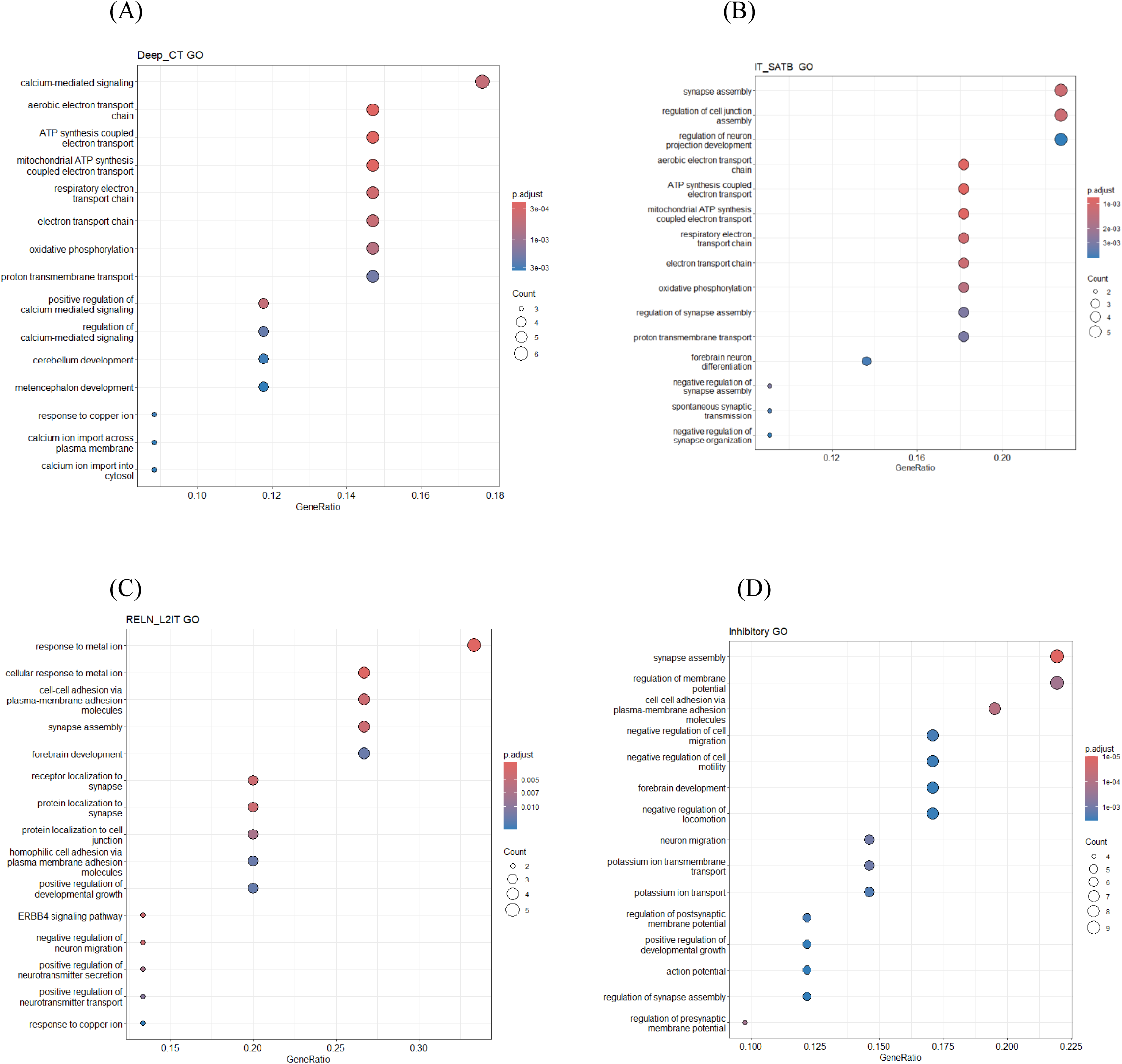
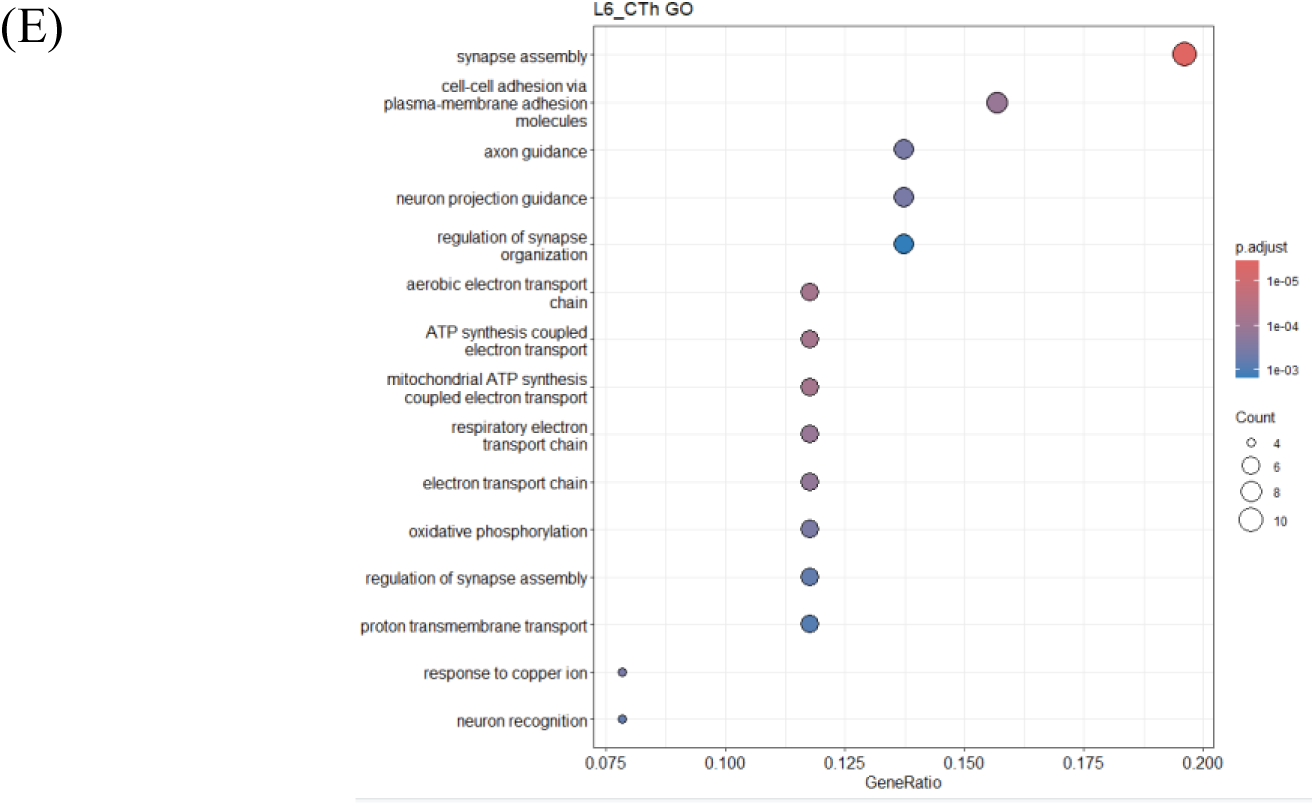
GO biological pathway analysis for enriched transcripts within (A) Deep_CT neuron, (B) IT_SATB neuron, (C) RELN neuron, (D) Inhibitory neuron in DLPFC, (E) L6_CTh neuron.

**S3.**
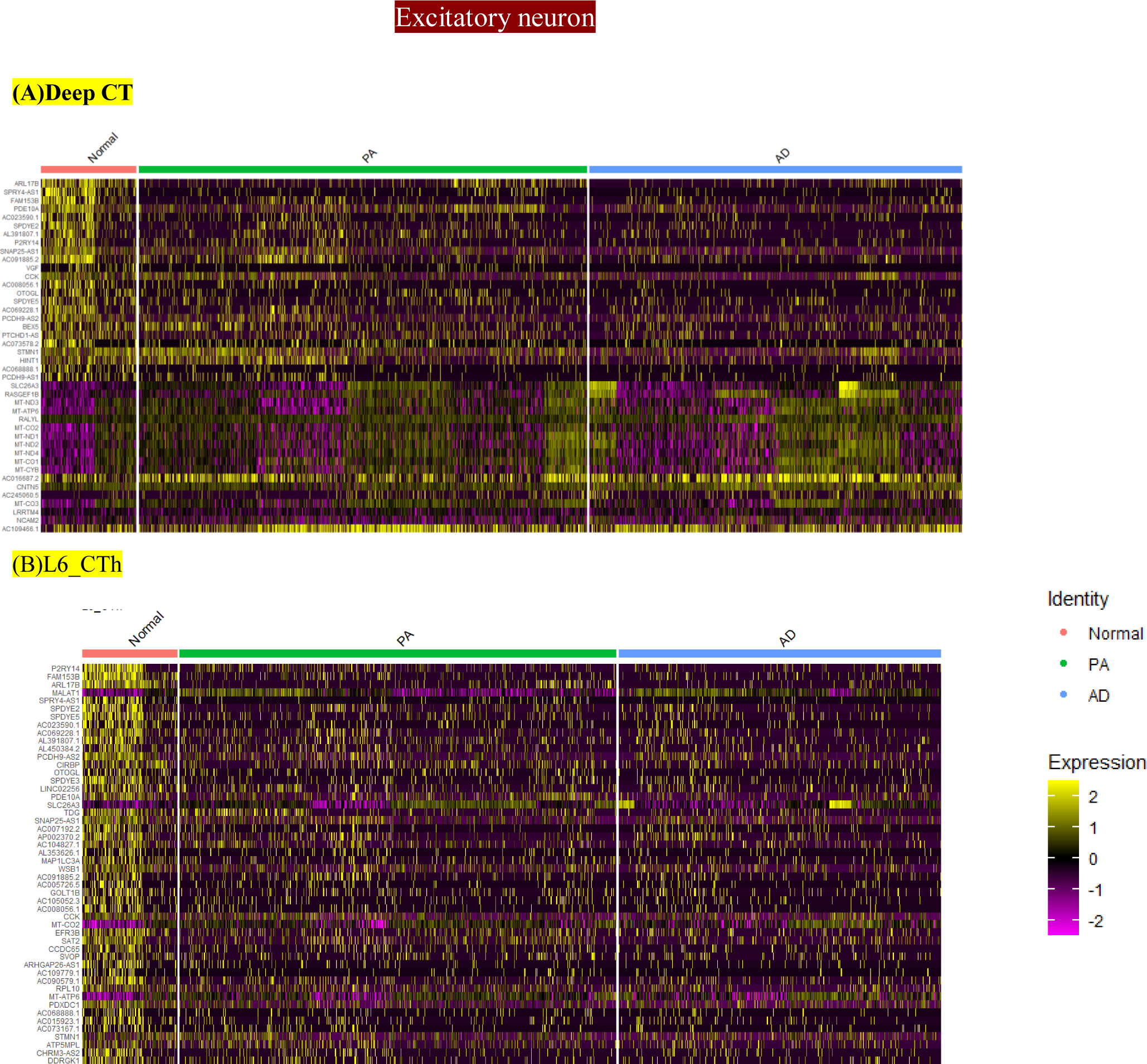
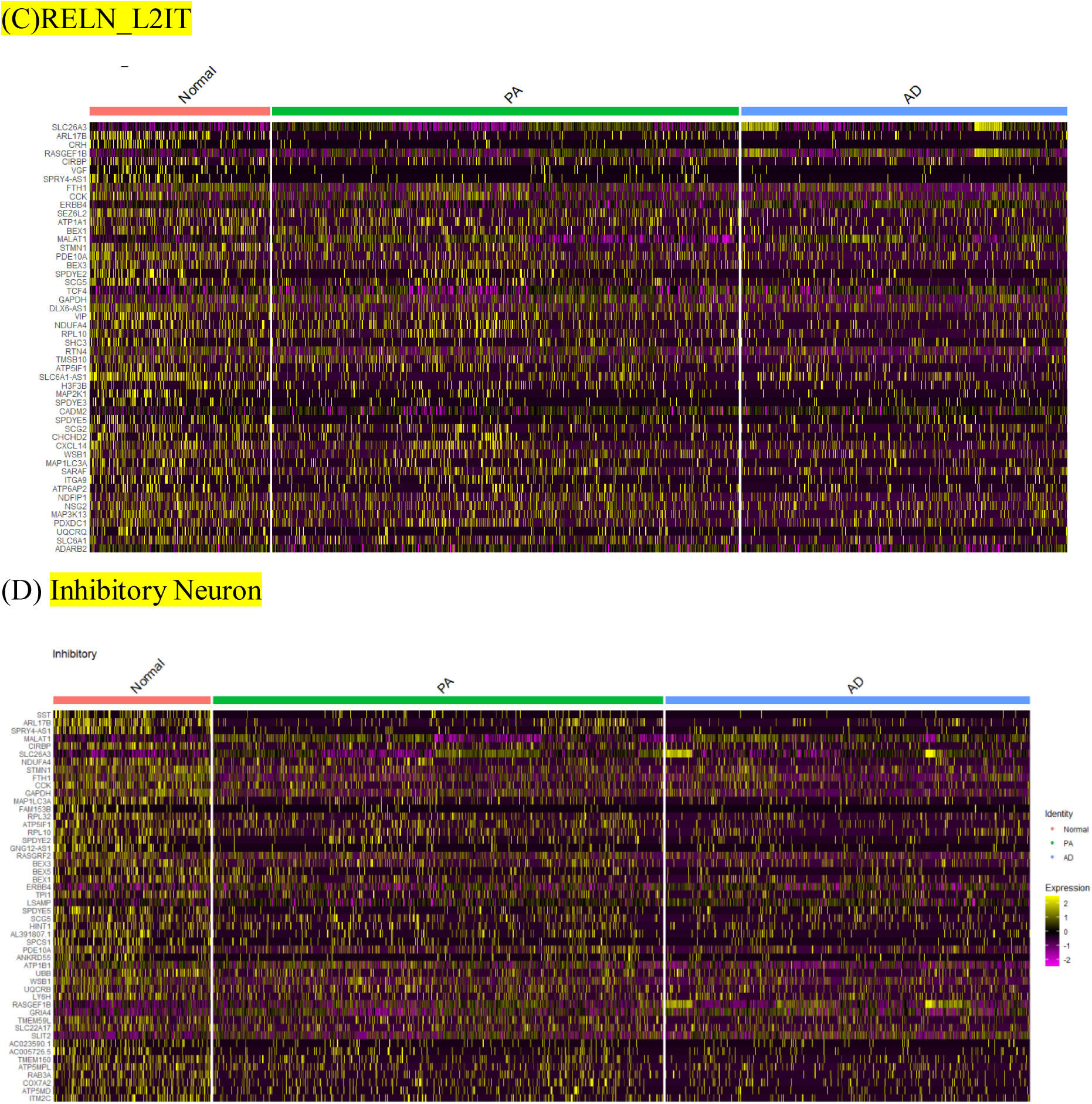
Heatmap denotes the expression pattern of top enriched genes in (A) Deep_CT, (B) L6_CTh, (C) RELN, (D) IT_SATB across conditions (CN, PA, AD) in DLPFC. Intensity is represented in terms of Z score, (D) Inhibitory Neuron.

**S4.**
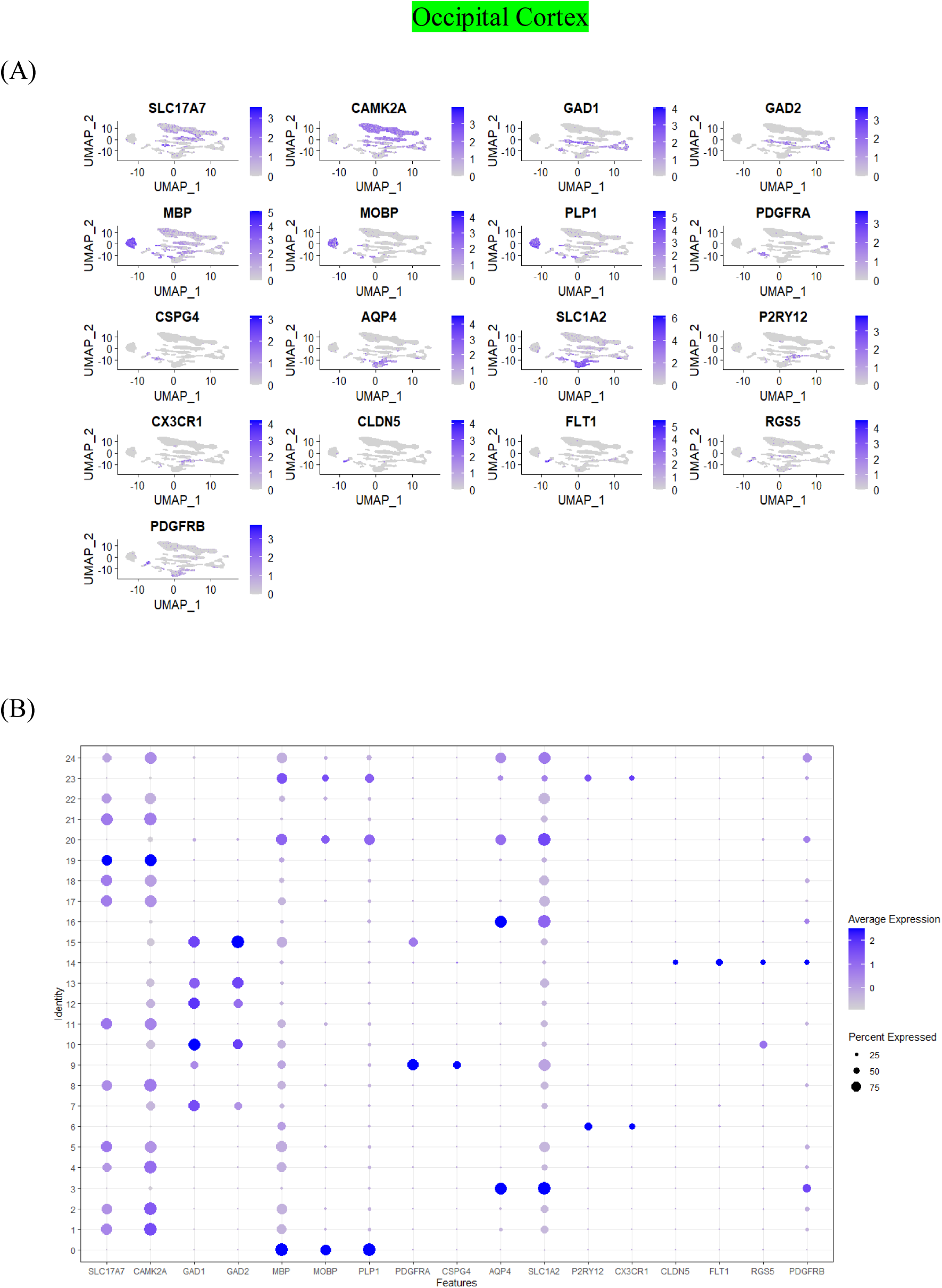
(A) Major Cell-type specific markers distribution in occipital cortex is denoted by UMAP. (B) Dot plots.

**S5.**
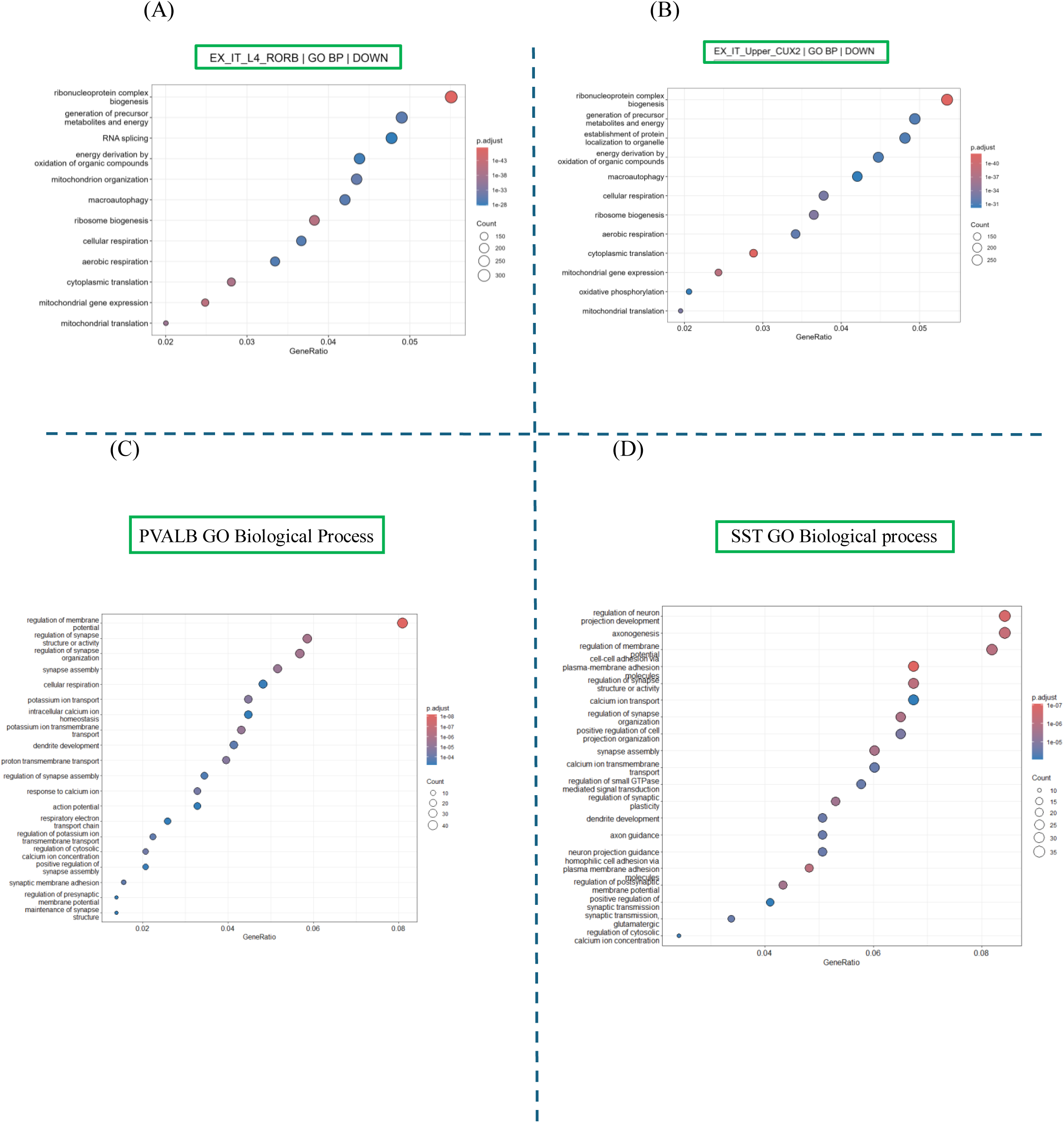
Go terms highlighted compromised biological pathways in (A) L4_RORB neuron and (B) CUX2 positive neuron; and Enriched biological pathways in Inhibitory neuron subpopulation: (C) PVALB positive Neuron. (D) SST neuron in the context of NHD in OC.

**S6.**
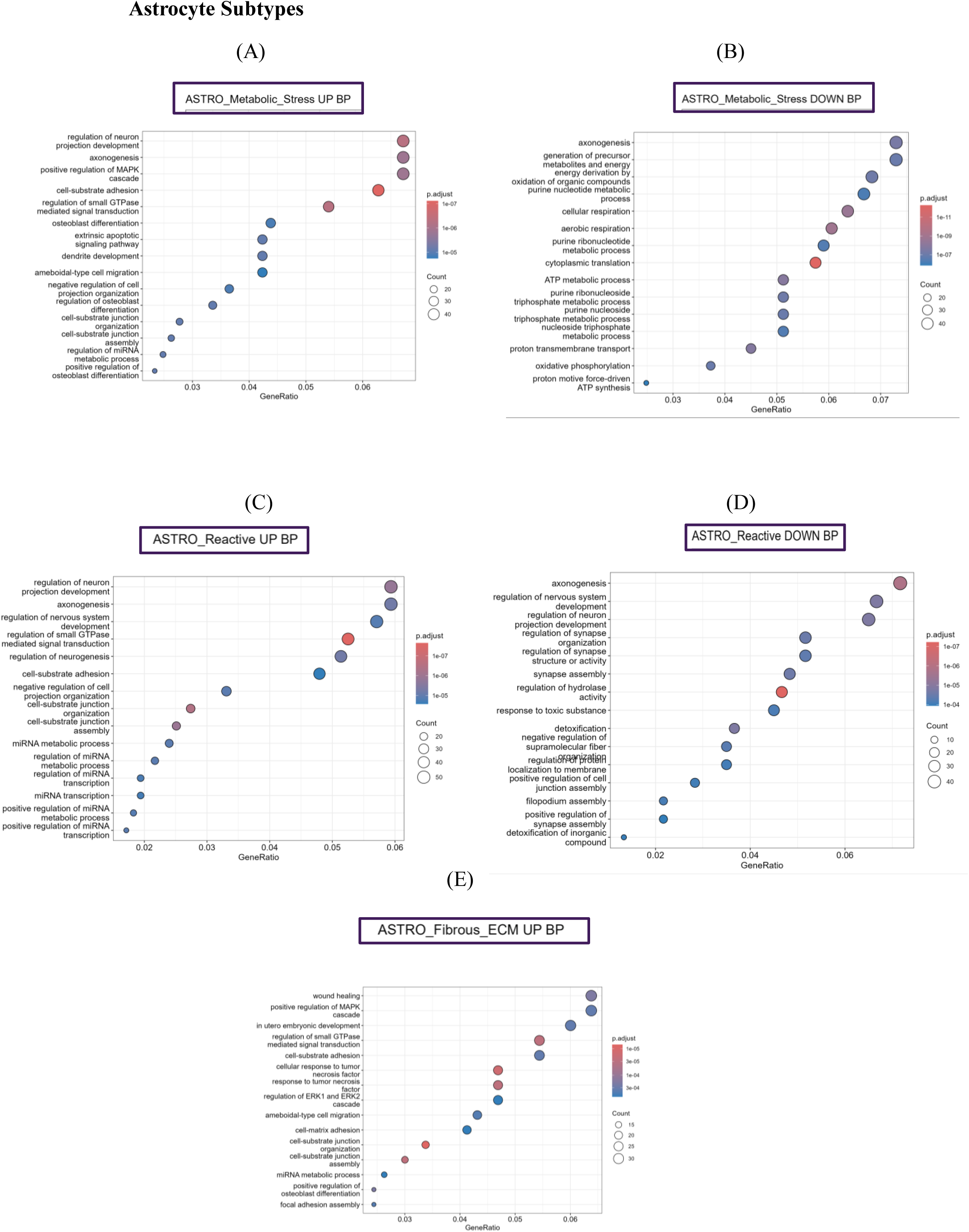
GO analysis highlighted Biological Pathways that are upregulated in (A) Metabolic stress associated astrocytes (Astro_Metabolic stress),(C) Reactive Astrocytes, (E) Fibrous Astrocytes and downregulated in (B) Metabolic stress associated astrocytes, (D) Reactive astrocytes.

**S7.**
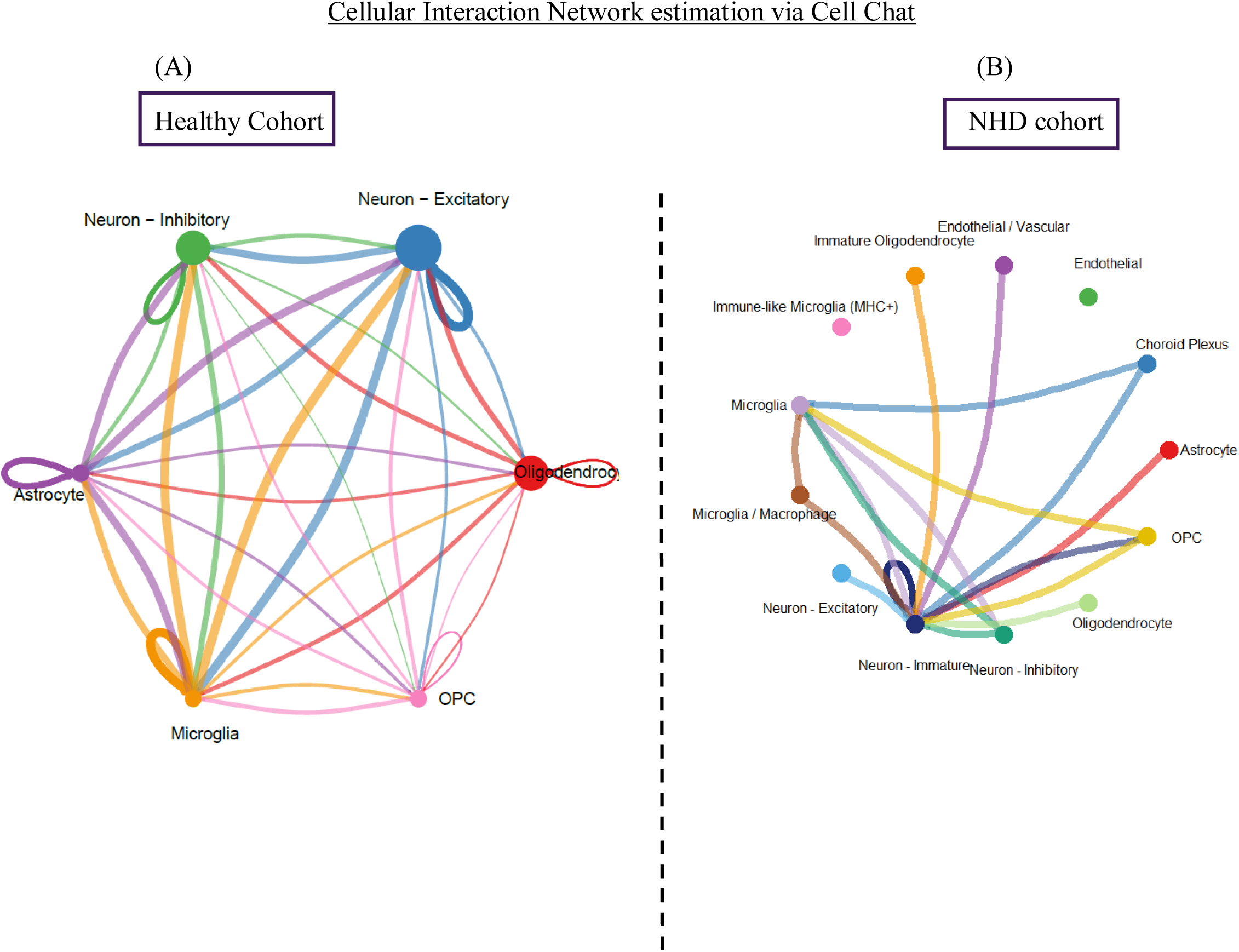
Cell-chat based estimation of cellular coordination network in OC: (A) In Healthy individuals we observed coherent interactions between major cell types, whereas in (B) NHD individual that pattern is collapsed and the coordination is majorly dominated by glial population.

**S8.**
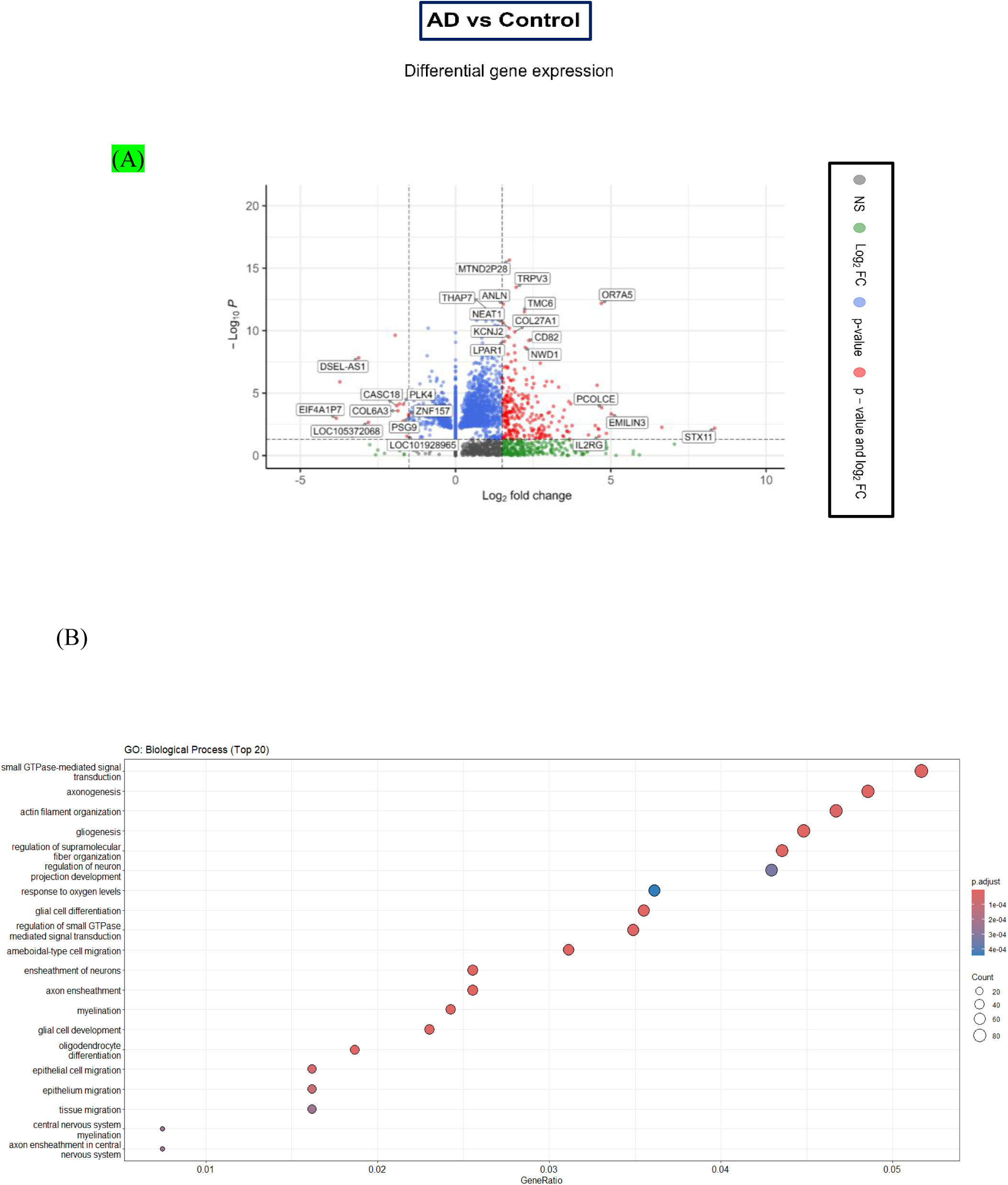
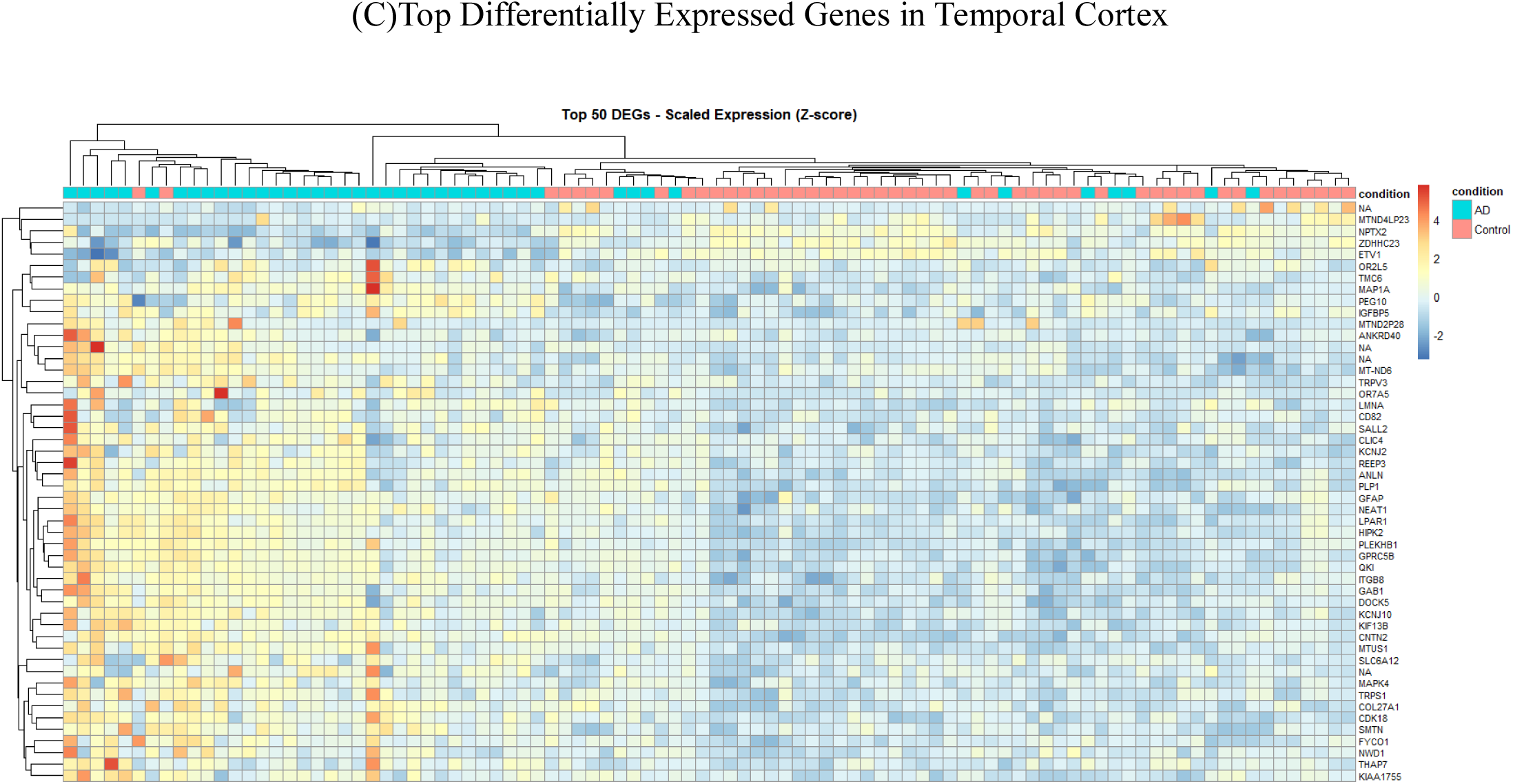
Temporal cortex. **(A)** volcano plot represents DEGs in Alzheimer’s brain as compared to Cognitive normal**. (B)** GO enrichment analysis highlights biological process associated with upregulated transcripts in Alzheimer’s panel. (C) Heatmap implies major gene expression patterns in AD vs Control brain.

**S9 :**
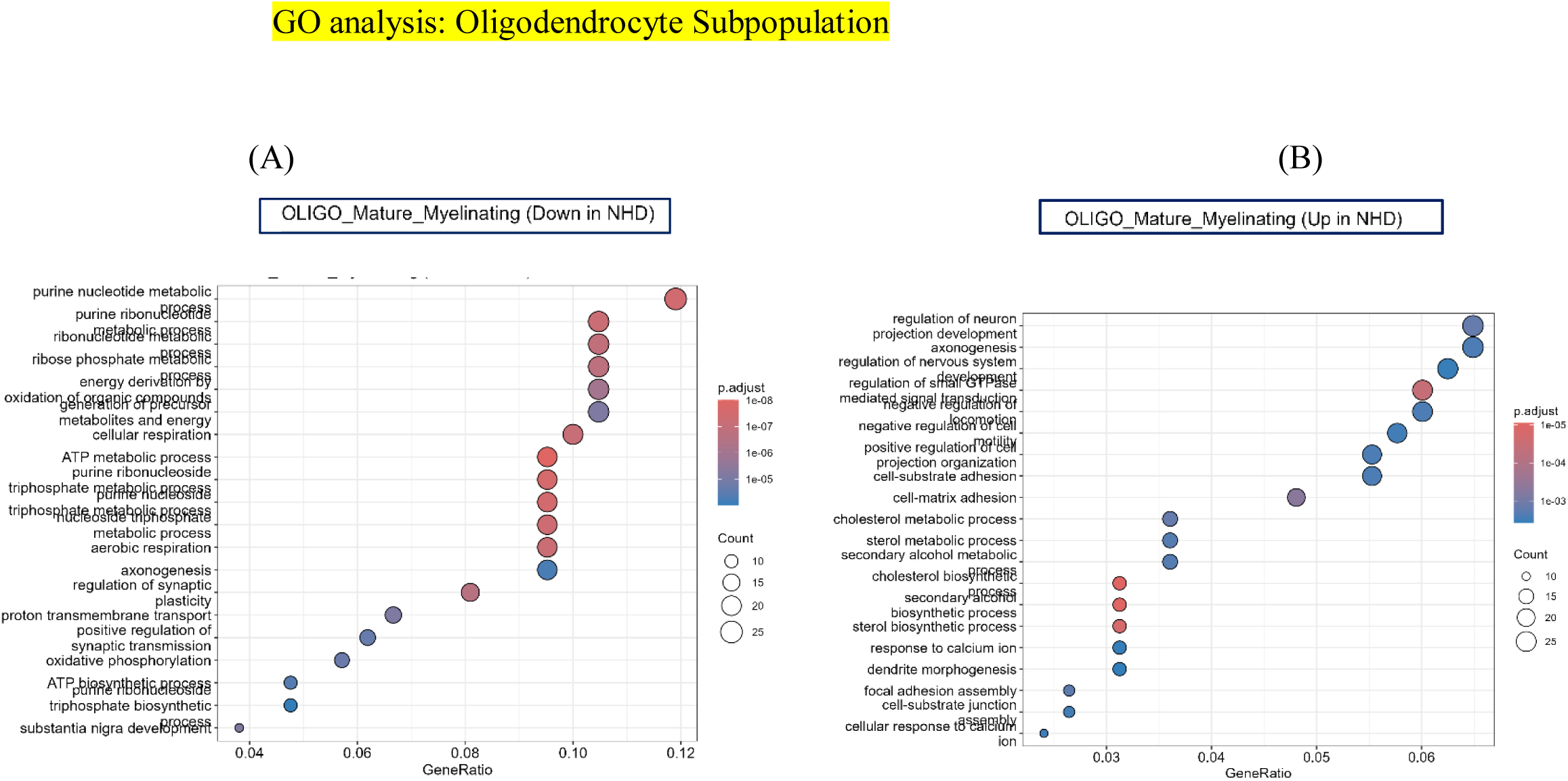
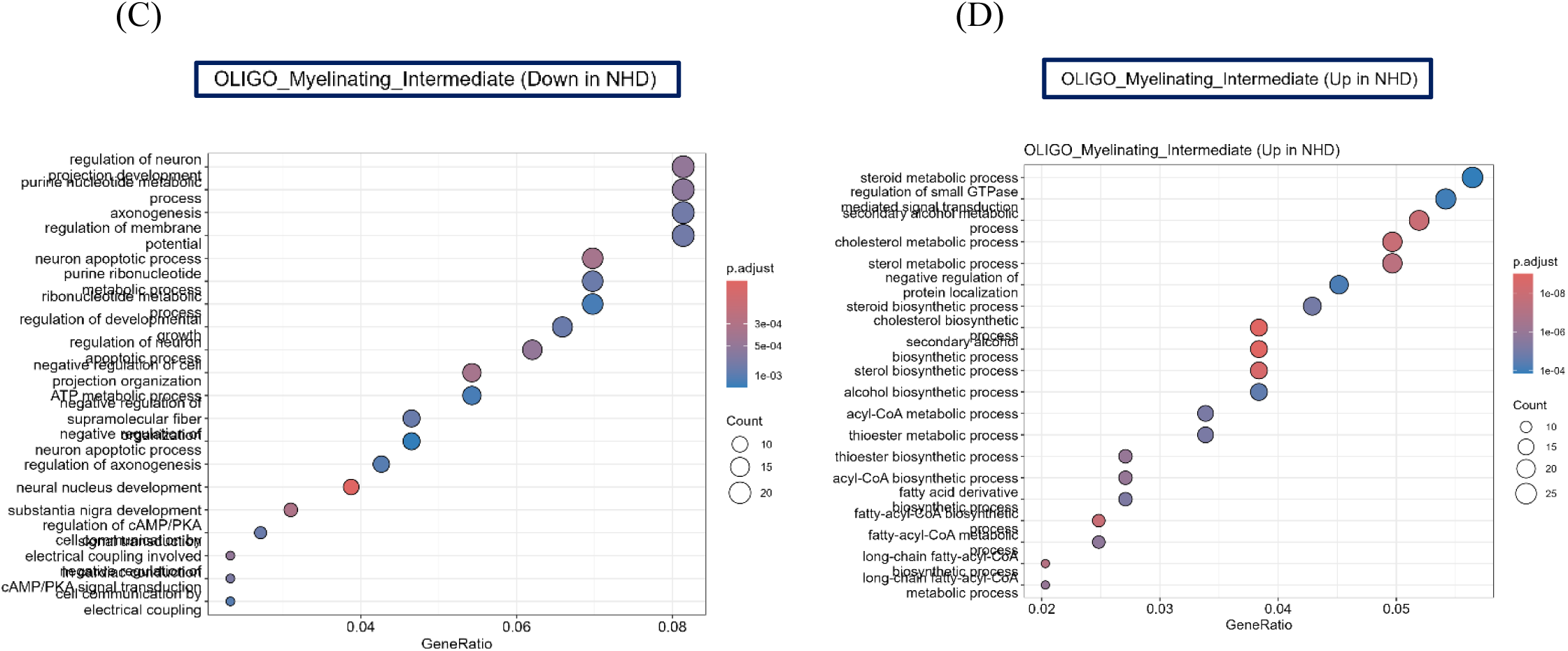
Oligodendrocyte subpopulation GO analysis (A) Mature myelinating upregulation in NHD (B) Mature myelinating downregulation in NHD; (C) Myelinating_Intermediate upregulation in NHD and (D) Myelinating_Intermediate downregulation in NHD.

